# Microglial depletion with CSF1R inhibitor during chronic phase of experimental traumatic brain injury reduces neurodegeneration and neurological deficits

**DOI:** 10.1101/791871

**Authors:** Rebecca J. Henry, Rodney M. Ritzel, James P. Barrett, Sarah J. Doran, Yun Jiao, Jennie B. Leach, Gregory L. Szeto, Bogdan A. Stoica, Alan I. Faden, David J. Loane

## Abstract

Chronic neuroinflammation with sustained microglial activation occurs following moderate-to-severe traumatic brain injury (TBI) and is believed to contribute to subsequent neurodegeneration and neurological deficits. Microglia, the primary innate immune cells in brain, are dependent on colony stimulating factor 1 receptor (CSF1R) signaling for their survival. In this translational study, we examined the effects of delayed depletion and subsequent repopulation of microglia on chronic neurodegeneration and functional recovery up to three months posttrauma. A CSF1R inhibitor, PLX5622, was administered to injured adult male C57Bl/6 mice at one month after controlled cortical impact to remove chronically activated microglia, and the inihibitor was withdrawn 1 week later to allow microglial repopulation. Following TBI, the repopulated microglia displayed a ramified morphology, similar to that of sham control uninjured animals, whereas microglia in untreated injured animals showed the typical chronic posttraumatic hypertrophic morphology. PLX5622 treatment limited TBI-associated neuropathological changes at 3 months posttrauma; these included a smaller cortical lesion, reduced neuronal cell death in the injured cortex and ipsilateral hippocampus, and decreased NOX2-dependent reactive microgliosis. Furthermore, delayed depletion of microglia led to widespread changes in the cortical transcriptome, including alterations in gene pathways involved in neuroinflammation, oxidative stress, and neuroplasticity. PLX5622 treated animals showed significantly improved neurological recovery using a variety of complementary neurobehavioral evaluations. These included beam walk and rotorod tests for sensori-motor function, as well as Y-maze, novel object recognition, and Morris water maze tests to evaluate cognitive function. Together, our findings show that chronic phase removal of neurotoxic microglia using CSF1R inhibitors after experimental TBI can markedly reduce chronic neuroinflammation and neurodegeneration, as well as related long-term motor and cognitive deficits. Thus, CSF1R inhibition may provide a clinically feasible approach to limit posttraumatic neurodegeneration and neurological dysfunction following head injury.

## Introduction

Traumatic brain injury (TBI) is a major cause of morbidity and mortality world-wide (Maas *et al*., 2017). Its high incidence and prevalence result in a large number of survivors with significant disabilities (Bryant *et al*., 2010; Sabaz *et al*., 2014; Kumar *et al*., 2018). Traumatic insults to the brain produce tissue damage and related neurological dysfunction through both direct mechanical damage (primary injury) and more delayed mechanisms that result from secondary biochemical/metabolic alterations (Loane and Byrnes, 2010; Faden and Loane, 2015). Recent clinical and experimental studies have implicated chronic neuroinflammation as an important secondary injury mechanism that may be treatable (Loane and Byrnes, 2010; Simon *et al*., 2017). Such neuroinflammation includes activation of brain resident microglia.

Microglia play a critical role in neuronal plasticity and maintaining tissue homeostasis in brain. Following brain trauma, microglia become activated and undergo both phenotypic and functional alterations. Microglial activation is an essential innate immune process that is required to protect the host from injury by promoting phagocytosis of dying or dead cells during recovery or repair from brain injury. However, uncontrolled activation, or activation over extended periods, may be detrimental, contributing to both subsequent neurodegenerative processes and related behavioral dysfunction (Block and Hong, 2005; Loane and Byrnes, 2010). Chronic microglial activation has been reported in TBI patients for many years following the initial brain trauma (Gentleman *et al*., 2004; Ramlackhansingh *et al*., 2011; Johnson *et al*., 2013; Coughlin *et al*., 2015; Cherry *et al*., 2016; Coughlin *et al*., 2017), and is more broadly distributed than suggested by the primary lesion. Moreover, it is implicated in persistent white matter atrophy and related cognitive impairments (Maxwell *et al*., 2006; Bendlin *et al*., 2008; Ramlackhansingh *et al*., 2011; Johnson *et al*., 2013; Coughlin *et al*., 2015; Wilson *et al*., 2017).

Microglia express colony stimulating factor 1 receptor (CSF1R) (Erblich *et al*., 2011; Nandi *et al*., 2012), which is essential for microglial maintenance and long-term survival (Ginhoux *et al*., 2010; Erblich *et al*., 2011). The recent development of CSF1R inhibitors has enabled specific targeting of microglia under both physiolgical (Elmore *et al*., 2014) and pathological states (Szalay *et al*., 2016; Jin *et al*., 2017; Li *et al*., 2017), with differential effects on function reported depending on the timing of microglia depletion. Oral administration of CSF1R inhibitors can cause rapid depletion of >95% of all microglia within 7 days of treatment (Elmore *et al*., 2014). In a mouse hippocampal lesion model, depletion of microglia by 30-day treatment with a CSF1R inhibitor, PLX3397, improved functional recovery and reduced pro-inflammatory gene expression in hippocampus (Rice *et al*., 2015). Despite evidence that chronic evolving microglial activation is detrimental following moderate-severe TBI (Byrnes *et al*., 2012; Loane *et al*., 2014; Pischiutta *et al*., 2018), prolonged inhibition of microglia is unlikely to be therapeutic because microglia perform critical functions such as synaptic pruning and restoration of tissue homeostasis after pathological insult (Salter and Stevens, 2017). However, withdrawal of CSF1R inhibitors in microglia-depleted mice resulted in rapid repopulation of microglia (Elmore *et al*., 2014; Elmore *et al*., 2015), which correlated with improved functional recovery in a hippocampal lesion model in mice (Rice *et al*., 2017). Therefore, removing pro-inflammatory and neurotoxic microglial populations by short-term depletion followed by repopulation may have a long-term effect on the inflammatory lesion environment and disrupt the chronic trajectory of neurodegeneration, leading to enhanced functional recovery after TBI.

Here, we examined whether pharmacological depletion and subsequent restoration of microglia long after moderate-severe TBI in a well-characterized mouse model, can reduce posttraumatic neurodegeneration and limit long-term neurological dysfunction. Our findings strongly implicate neurotoxic microglial activation as a major pathophysiological factor in chronic posttraumatic neurodegeneration and related neurological dysfunction after brain trauma, and suggest a new potential therapeutic target for this disorder.

## Materials and Methods

### Animals

Studies were performed using adult male C57Bl/6J mice (10-12 weeks old; Taconic Biosciences Inc., Germantown, NY). Mice were housed in the Animal Care facility at the University of Maryland School of Medicine under a 12-hour light-dark cycle, with *ad libitum* access to food and water. All surgical procedures were carried out in accordance with protocols approved by the Institutional Animal Care and Use Committee (IACUC) at the University of Maryland School Of Medicine.

### Controlled cortical impact

Our custom-designed CCI injury device consists of a microprocessor-controlled pneumatic impactor with a 3.5 mm diameter tip. Briefly, mice were anesthetized with isoflurane evaporated in a gas mixture containing 70% N_2_O and 30% O_2_ and administered through a nose mask (induction at 4% and maintenance at 2%). A 10 mm midline incision was made over the skull, the skin and fascia were reflected, and a 4 mm craniotomy was made on the central aspect of the left parietal bone. The impounder tip of the injury device was then extended to its full stroke distance (44 mm), positioned to the surface of the exposed dura, and reset to impact the cortical surface. Moderate-level CCI was induced using an impactor velocity of 6 m/s and deformation depth of 2 mm (Loane *et al*., 2014; Kumar *et al*., 2016a; Henry *et al*., 2019). Sham animals underwent the same procedure as TBI mice except for the craniotomy and impact.

### Plexxikon (PLX) 5622 administration

PLX5622 was provided by Plexxikon Inc. (Berkley, CA) and formulated in AIN-76A rodent chow by Research Diets Inc. (New Brunswick, NJ) at a concentration of 1200 ppm (Spangenberg *et al*., 2019). Standard AIN-76A diet was provided as vehicle control. Mice were provided ad libitum acess to PLX5622 or vehicle diet for 7 days (d) to deplete microglia. This dose and time was valid to deplete 95% of microglia (**Fig S1A, S1B**).

### Experimental design

C57Bl/6 mice (n=15-20/group) were subjected to either controlled cortical injury (CCI) or Sham surgery. At 28 days post-injury (dpi), animals were placed on PLX5622 (1200 ppm; Plexxikon Inc.) or Vehicle chow for one week to deplete microglia (**Fig. S2**). Animals were returned to a standard chow diet to allow for microglial repopulation, and were maintained on standard chow for the remainder of the study. Following completion of comprehensive behavioural testing at 84 dpi, mice were anesthetized (100 mg/kg sodium pentobarbital, I.P.) and transcardially perfused with ice-cold 0.9% saline (100ml), followed by 300 ml of 4% paraformaldehyde (PFA). Brains were removed and post-fixed in 4% PFA overnight, cryoprotected in 30% sucrose, and were processed for histological outcome measures. A separate cohort of mice (n=8-9/group) underwent either CCI or Sham surgery. At 28 dpi, animals were placed on PLX5622 or control chow as described above. At 56 dpi, mice were anesthetized (100 mg/kg sodium pentobarbital, I.P.) and transcardially perfused with ice-cold 0.9% saline (100ml), the ipsilateral brain tissue was rapidly dissected and processed for flow cytometry, or RNA/Nanostring analysis.

### Neurobehavioral testing

#### Beam walk

Fine motor coordination was assessed using a beam walk test, as previously described (Kumar *et al*., 2016b; Henry *et al*., 2019). Briefly, mice were placed on one end of a wooden beam (5 mm wide and 120 mm in length), and the number of foot faults of the right hind limb were recorded over 50 steps. Mice were trained on the beam walk for 3 d prior to Sham or CCI and tested weekly through 84 dpi.

#### Accelerating rotorod

Gross motor function and balance was assessed on the accelerating Rotorod at 84 dpi, as previously described (Doran *et al*., 2019). On the day of testing, mice were placed on the rod which accelerated from 4-60 RPM with maximal speed reached within 20 sec. Latency to fall from the rod (or cling to and rotate with the rod for three consecutive rotations) was recorded for three trials. A 5–10 min of rest period with access to food and water between was allowed between each trial. Scores from the three trials were averaged to give a single score for each mouse.

#### Y-maze maze

The Y-maze was performed at 70 dpi to test spatial working memory in mice, as previously described (Kumar *et al*., 2016b; Henry *et al*., 2019). Briefly, the Y-maze (Stoelting Co., Wood Dale, IL) consisted of three identical arms, each arm 35 cm long, 5 cm wide, and 10 cm high, at an angle of 120° with respect to the other arms. One arm was randomly selected as the “start” arm, and the mouse was placed within and allowed to explore the maze freely for 5 min. Arm entries (arms A–C) were recorded by analyzing mouse activity using ANY-maze software (Stoelting Co., Wood Dale, IL). An arm entry was attributed when all four paws of the mouse entered the arm, and an alternation was designated when the mouse entered three different arms consecutively. The percentage of alternation was calculated as follows: total alternations x 100/(total arm entries − 2). If a mouse scored significantly >50% alternations (the chance level for choosing the unfamiliar arm), this was indicative of spatial working memory.

#### Novel object recognition (NOR)

NOR was performed on 77-78 dpi to assess non-spatial hippocampal-mediated memory, as previously described (Zhao *et al*., 2012). Mice were placed in an open field (22.5 cm X 22.5 cm) and two identical objects were placed near the left and right corners of the open field for training (familiar phase). Mice were allowed to freely explore until they spent a total of 20 sec exploring the objects (exploration was recorded when the front paws or nose contacted the object). The time spent with each object was recorded using Any-Maze software (Stoelting Co.). After 24 h, object recognition was tested by substituting a novel object for a familiar training object (the novel object location was counterbalanced across mice). Because mice inherently prefer to explore novel objects, a preference for the novel object (more time than chance [15 sec] spent with the novel object) indicates intact memory for the familiar object.

#### Morris water maze (MWM)

Spatial learning and memory was assessed using the MWM, as previously described (Byrnes *et al*., 2012; Zhao *et al*., 2012). The MWM protocol included two phases: (1) standard hidden platform training (learning acquisition) and (2) a twenty-four-hour probe test (reference memory). Briefly, a circular tank (100 cm in diameter) was filled with water (23±2°C) and was surrounded by various extra-maze cues on the wall of the testing area. A transparent platform (10 cm in diameter) was submerged 0.5 cm below the surface of the water. Starting at 80 dpi, the mice were trained to find the hidden submerged platform located in the northeast (NE) quadrant of tank for 4 consecutive days (80-84 dpi). The mice underwent four trials per day, starting from a randomly-selected release point (east, south, west, and north). Each mouse was allowed a maximum of 90 sec to find the hidden submerged platform. The latency to find the submerged platform was recorded using Any-Maze software (Stoelting Co.). Reference memory was assessed by a probe test carried out 24 h after the final acquisition day. The platform was removed and the mice were released from the southwest (SW) position, and the time in the target quadrant was recorded. Search strategy analysis was performed, as previously described (Byrnes *et al*., 2012; Zhao *et al*., 2012). Three strategies were identified using the following categorization scheme (**Fig. S5B**): a) *spatial search strategy* was defined as swimming directly to platform with no more than 1 loop, or swimming directly to the correct target quadrant and searching for platform; b) *non-spatial systematic strategy* was defined as searching interior portion of or entire tank without spatial bias, including searching within incorrect target quadrant prior to finding platform; c) *repetitive looping strategy* was defined as circular swimming around the tank, swimming in tight circle, or swimming around the wall of tank before finding the submerged hidden platform. The search strategies were analyzed during the MWM probe test, and percentage of each strategy in each group was calculated using a chi-square analysis. Mice that remained immobile throughout the 90 sec trial in the MWM test were excluded from the analysis (Sham: n=2; TBI+Veh: n=3; TBI+PLX: n=2).

### Histology

#### Lesion volume

Lesion volume was measured on 60 μm coronal sections that were stained with cresyl violet (FD NeuroTechnologies, Baltimore, MD; n=8-12/group). Quantification was based on the Cavalieri method using Stereoinvestigator Software (MBF Biosciences, Williston, VT), as previously described (Byrnes *et al*., 2012; Henry *et al*., 2019). Briefly, the lesion volume was quantified by outlining the missing tissue on the injured hemisphere using the Cavalieri estimator with a grid spacing of 0.1 mm. Every 8^th^ section from a total of 96 sections was analyzed beginning from a random start point.

#### Neuronal loss

For analysis of post-traumatic neuronal cell loss 60µm coronal sections were stained with cresyl violet, and the optical fractionator method of stereology using Stereoinvestigator Software (MBF Biosciences) was employed, as previously described (Loane *et al*., 2014). Neurons in the ipsilateral cortex were characterized according to previously described methods (Garcia-Cabezas *et al*., 2016). For neuronal loss in cortex and dentate gyrus (DG) region of the hippocampus, every fourth 60 µm section between −1.34 and −2.54 mm and −2.70 to −3.16 mm, respectively, from bregma was analyzed beginning from a random start point. A total of 6 sections were analyzed. The number of surviving neurons in each field was divided by the volume of the region of interest to obtain the cellular density expressed in counts/mm^3^.

#### Microglia morphological analysis

Immunohistochemistry was performed on 60 μm coronal sections to assess morphological activation states of microglia in the perilesional cortex, as previously described (Byrnes *et al*., 2012; Loane *et al*., 2014). Briefly, sections were incubated overnight with rabbit anti-Iba-1 (1:5000; Wako Chemicals, Richmond, VA), washed in 1x PBS (three times), and incubated with biotinylated anti-rabbit IgG antibody (Vector Laboratories, Burlingame, CA) for 2 h at room temperature. Sections were incubated in avidin-biotin-horseradish peroxidase solution (Vectastain Elite ABC kit; Vector Laboratories) for 1 hour and then reacted with 3,3¢-diaminobenzidine (Vector Laboratories) for color development. Iba-1 stained sections were counterstained with Cresyl violet and mounted for immunohistochemical analysis using a Leica DM4000B microscope (Leica Microsystems; Exton, PA). Stereoinvestigator Software (MBF Biosciences) was used to count and classify the number of cortical microglia for microglial morphologic phenotypes (ramified and hypertrophic/bushy) at 84 dpi, using the optical fractionator method (n=10-11/group).

#### Immunofluorescence imaging

Immunofluorescence imaging was carried out on 20 µm coronal brain sections at approximately −1.70 mm from bregma. Standard immunostaining techniques were employed, as previously described (Henry *et al*., 2019). Briefly, 20 µm brain sections were washed three times with 1x PBS, blocked for 1 hour in goat serum containing 0.4% Triton X-100, and incubated overnight at 4°C with a combination of primary antibodies, including mouse anti-gp91^phox^ (NOX2; 1:1000, BD Biosciences, San Jose, CA), rat anti-CD68 (1:1000, AbD Serotec Inc., Raleigh, NC), rabbit anti-Iba1 (1:1000, Wako Chemicals) and chicken anti-GFAP (1:500, Abcam, Eugene, OR). Sections were washed with 1x PBS (three times), and incubated with appropriate Alexa Fluor conjugated secondary antibodies (Life Technologies, Carlsbad, CA) for 2 h at room temperature. Sections were washed with 1x PBS (three times), counterstained with 4’, 6-diamidino-2-phenylindole (DAPI; 1 μg/ml; Sigma, Dorset, UK), and mounted with glass coverslips using Hydromount solution (National diagnostics, Atlanta, GA). Images were acquired using a fluorescent Nikon Ti-E inverted microscope (Nikon Instrument Inc., Melville, NY), at 10x (Plan Apo 10X NA 0.45) or 20x (Plan APO 20X NA 0.75) magnification. Exposure times were kept constant for all sections in each experiment. All images were quantified using Nikon ND-Elements Software (AR 4.20.01). Co-localization of NOX2 and CD68 in microglia/macrophage (Iba1+) was performed by binary operation intersection followed by thresholding. 12 positive regions of interest near the lesion site per mouse were quantified (with n=5 mice/group), and was expressed as NOX2+CD68+ cells/mm^2^. For GFAP quantification, tiled images of each brain section were at 10x and 20x magnification (n=5-8 mice/group). The background was subtracted using a specific reference region just outside each brain section. The quantified areas were the ipsilateral perilesional region (proximal to the lesion site) and the corresponding site in sham animals. The total GFAP immunofluorescence in the perilesional region was recorded and presented following normalization to the perilesional region area. A common threshold was selected for all images to identify the specific cellular (astrocytic) GFAP intensity. The area occupied by astrocytes in the perilesional region as well as the GFAP immunofluorescence in the astrocytic fraction was quantified and presented following normalization to the area occupied by astrocytes in the perilesional region.

### Molecular and Cellular analysis

#### Nanostring analysis

RNA was extracted and purified from frozen cortical tissue using an RNA Plus Universal Mini Kit (Qiagen, Hilden, Germany). Total RNA was diluted to 20 ng/μl and probed using an nCounter^©^ Mouse Neuropathology panel (Nanostring Technologies, Seattle, WA) profiling 770 genes across six fundamental themes of neurodegeneration: neurotransmission, neuron-glia interaction, neuroplasticity, cell structure integrity, neuroinflammation, and metabolism. Counts for target genes were normalized to the best fitting house-keeping genes as determined by nSolver software.

Partial Least Squares Discriminant Analysis (PLSDA): PLSDA was performed with mixOmics v6.8.2 (Rohart *et al*., 2017) using normalized gene counts. All discriminant models were evaluated based on ROC curve using 100 iteration of cross-validation and bootstrap resampling with ROC632 v0.6 (Foucher and Danger, 2012). Pairwise differential expression analyses were performed with NanoStringDiff (v3.6.0) (Wang *et al*., 2016) using raw gene counts along with positive and negative controls and housekeeping genes from NanoString nCounter. Four comparisons were performed: 1) Sham+Veh vs Sham+PLX, 2) Sham+Veh vs TBI+Veh, 3) TBI+Veh vs TBI+PLX, and 4) TBI+PLX vs Sham+PLX. Differentially expressed genes were defined as those with an adjusted p-value <0.1 for either comparison 1 or 3. Results were plotted using p.heatmap v1.0.12 with hierarchical clustering of genes and experimental groups for all differentially expressed genes. Gene expression was normalized across groups as z-scores. Heatmaps were also independently plotted and clustered by genes only on annotated subsets with defined functional classes. MixOmics, ROC632, NanoStringDiff, and Pheatmap packages were all used in Rstudio v3.6.0. Violin plots of normalized gene counts were generated using Prism v8.0.2 (Graphpad), and annotated by median and upper/lower quartiles. Corresponding negative log10 adjusted p-values were plotted for the pairwise comparisons shown as calculated by NanoStringDiff.

#### Real-Time PCR

Quantitative gene expression analysis in the cortex was performed using Taqman technology, as previously described (Barrett *et al*., 2017; Henry *et al*., 2019). Target mRNAs included TaqMan gene expression assays for *Cybb* (NOX2) Mm01287743_m1; *Cyba* (p22^phox^) Mm00514478_m1; *Ncf1* (p47^phox^) Mm00447921_m1; *Ncf4* (p40^phox^) Mm00476300_m1; *Il1b* (IL-1β) Mm01336189_m1; *Nlrp3* (NLRP3) Mm00840904_m1; *Casp1* (Caspase-1) Mm00438023_m1; *Gfap* (GFAP) Mm01253033_m1; *Gapdh* (GAPDH) Mm99999915_g1; Applied Biosystems, Carlsbad, CA), and analysis was performed on ABI 7900 HT FAST Real Time PCR machine (Applied Biosystems). Samples were assayed in duplicate in one run (40 cycles), which was composed of 3 stages: 50°C for 2 min, 95°C for 10 sec for each cycle (denaturation), and a transcription step at 60°C for 1 min. Gene expression was normalized by GAPDH and compared to the control sample to determine relative expression values by 2^−ΔΔ^*^Ct^* method.

#### Flow cytometry

The ipsilateral hemisphere of each mouse was placed in complete Roswell Park Memorial Institute (RPMI) 1640 (Lonza Group, Basel, Switzerland) medium. Brain tissue was mechanically digested using a razor blade to mince tissue and was passed through a 70-μm-filter using RPMI. CNS tissue was then enzymatically digested using DNAse (10 mg/mL; Roche, Mannheim, Germany), Collagenase/Dispase (1 mg/mL; Roche), and Papain (25 U; Worthington Biochemical, Lakewood, NJ) for 1 h at 37°C in a shaking CO_2_ incubator (200 rpm). Tissue homogenates were centrifuged at 1500 rpm for 5 min at 4°C. The supernatant was discarded and the cells were resuspended in 70% Percoll^TM^ (GE Healthcare, Pittsburgh, PA) and underlayed in 30% Percoll^TM^. This gradient was centrifuged at 500 g for 20 min at 21°C. Myelin was removed by suction and cells at the interface were collected. Cells were washed and blocked with mouse Fc Block (Biolegend, San Diego, CA) prior to staining with primary antibody-conjugated flourophores: CD45-eF450, CD11b-APCeF780, Ly6C-APC (all Biolegend). For live/dead discrimination, a fixable viability dye, Zombie Aqua (Biolegend), was diluted at 1:100 in Hank’s balanced salt solution (HBSS; Gibco). Cells were briefly fixed in 2 % paraformaldehyde (PFA). Data were acquired on a LSRII using FACSDiva 6.0 (BD Biosciences, San Jose, CA) and analyzed using FlowJo (Treestar Inc., San Carlos, CA). The entirety of each sample was acquired on medium flow rate until zero events were obtained, providing absolute cell counts for each hemisphere. Resident microglia were identified as the CD45^int^CD11b^+^Ly6C^−^ population (Ritzel *et al*., 2019), whereas peripherally-derived immune cells were identified as CD45^hi^ as previously described (Ritzel *et al*., 2018b). Tissue and cell type matched fluorescence minus one (FMO) controls were used to determine the gating for each antibody. Prior to assessment on the cytometer, isolated cells were briefly probed to determine caspase-1 activity and IL-1β production. Immediately following percoll separation, cells were washed with PBS, and incubated with 1X FLICA solution at 37°C for 30 min according to the manufacturer’s instructions (ImmunoChemistry Technologies, Bloomington, MN). Cells were washed in FACS buffer, blocked, stained, and fixed as above. This assay employs the fluorescent inhibitor probe FAM-YVAD-FMK to label active caspase-1 enzyme in living cells (Lage *et al*., 2019). For Intracellular cytokine production analysis, prior to staining 1μl of GolgiPlug containing brefeldin A (BD Biosciences) was added to 800μl complete RPMI, and cells were incubated for 2 hours at 37°C (5% CO_2_). Cells were resuspended in Fc Block, stained for surface antigens, and washed in HBSS. Cells were vortexed and resuspended in 100μl of fixation/permeabilization solution (BD Biosciences) for 20 min. Cells were then washed twice in 300μl permeabilization/wash buffer (BD Biosciences), re-suspended in IL-1β-PerCPeF710 (Biolegend) for 30 min, washed, and subsequently fixed, as previously described (Ritzel *et al*., 2018a).

#### Statistical analysis

Randomization and blinding was performed as follows: a) individual who administered drugs was blinded to treatment group, and b) behavioral and stereological analyses were performed by individuals blinded to injury or treatment groups. Quantitative data were expressed as mean ± standard errors of the mean (s.e.m.). Normality testing was performed and data sets passed normality (D’Agostino & Pearson omnibus normality test), and therefore parametric statistical analysis was performed. Beam walk and acquisition days of the MWM was analyzed by one-way repeated measures ANOVA to determine the interactions of time and groups, followed by *post-hoc* adjustments using a Bonferroni’s multiple comparison test. Y maze, NOR, Rotorod, the MWM Probe trial, qRT-PCR, flow cytometry, microglia morphology and neuronal cell loss was analyzed by one-way ANOVA, followed by *post-hoc* adjustments using a Bonferroni’s multiple comparison test. Stereological data examining lesion volume and immunofluorescence was analyzed using a Student *t* test. The MWM search strategy was analyzed using a chi-square analysis. Statistical analyses were performed using GraphPad Prism Program, Version 8 for Windows (GraphPad Software, San Diego, CA, USA). A p<0.05 was considered statistically significant.

#### Data availability

The data used in this work will be available upon request. Nanostring data will be made available through GEOarchive.

## Results

### Delayed depletion of microglia with CSF1R inhibitor after TBI improves long-term motor and cognitive recovery

Microglia are critically dependent on CSF1R signaling for their survival (Elmore *et al*., 2014). PLX5622 is an orally bioavailable, brain-penetrant CSF1R inhibitor, that is able to achieve robust brain-wide microglia elimination (Spangenberg *et al*., 2019). To demonstrate its utility in our pre-clinical study, PLX5622 (1200 ppm) was formulated in rodent chow, and administered to adult male C57Bl/6J mice for 7 days. When compared to control levels in vehicle-treated mice, oral PLX5622 treatment led to almost complete microglial elimination (95% reduction) within 7 days (**Fig. S1A, S1B**). Returning PLX5622-treated mice to normal chow resulted in microglial repopulation in the brain within 7 days, such that numbers of repopulated microglia were not different to levels in control brain (**Fig. S1B**).

To investigate the effect of temporary depletion of microglia during the chronic stages of TBI on long-term neurological recovery, sham and moderate-level CCI mice were orally administered PLX5622 (1200 ppm) or Vehicle (oil) in chow for 1 week starting at 4 weeks post-injury. All mice were returned to normal chow at 5 weeks post-injury to allow for microglial repopulation, and comprehensive motor and cognitive functional testing was performed from 8 to 12 weeks (**Fig. S2**). Microglial depletion and repopulation do not negatively affect animal behavior in non-injured mice, including tests of anxiety, motor-function, and cognition (Rice *et al*., 2015; Spangenberg *et al*., 2019). Neurobehavioral testing confirmed that Sham+Vehicle and Sham+PLX6522-treated C57Bl/6J mice had equal performance in beam walk, Y-maze, novel object recognition and Morris water maze tests (**Fig. S3**). Therefore, Vehicle- and PLX5622-treated sham mice were combined (denoted sham) for all behavioral analyses.

TBI induces long-term deficits in fine motor coordination in a beam walk task. Prior to microglia depletion at 4 weeks post-injury, TBI mice had 42.8 ± 2.0 footfaults (ff; mean ± s.e.m.) when compared to sham mice (Sham = 7.1 ± 1.4 ff; *p*<0.001vs Sham; **Fig. 1A**). Sham and TBI mice were then randomized into cohorts that received PLX5622 or Vehicle for 1 week to deplete/repopulate microglia, and mice were tested weekly through the end of study. TBI-induced deficits in motor coordination persisted in Vehicle-treated TBI mice at 12 weeks post-injury (TBI+Veh = 45.7 ± 1.4 ff vs Sham = 10.6 ± 1.5 ff; TBI effect, F_(2,611)_=633.4, *p*<0.001; *p*<0.001 vs Sham; **Fig. 1A**). PLX5622 treatment significantly improved post-injury beam walk performance (F_(24,611)_=6.93, *p*<0.001). When compared to deficits in Vehicle-treated TBI mice, TBI+PLX5562-treated mice had reduced footfaults at 8 (TBI+Veh = 43.0 ± 3.1 ff vs TBI+PLX = 30.6 ± 4.4 ff; *p*<0.05), 10 (TBI+Veh = 44.6 ± 1.6 ff vs TBI+PLX = 28.8 ± 5.1 ff; *p*<0.05), and 12 (TBI+Veh = 45.7 ± 1.4 ff vs TBI+PLX = 30.7 ± 4.7; *p*<0.05) weeks post-injury. Motor balance was also evaluated using an accelerating rotarod task at 12 weeks post-injury. Vehicle-treated TBI mice spent less time on the accelerating rotorod than sham mice (TBI+Veh = 57.0 ± 7.2 sec vs Sham = 76.9 ± 3.2 sec; **Fig. S4**). In contrast, PLX5622-treated TBI mice spent more time on the rotarod (TBI+PLX = 70.3 ± 10.6 sec) than Vehicle-treated TBI mice, but differences did not reach statistical significance (**Fig. S4**).

**Figure 1.**
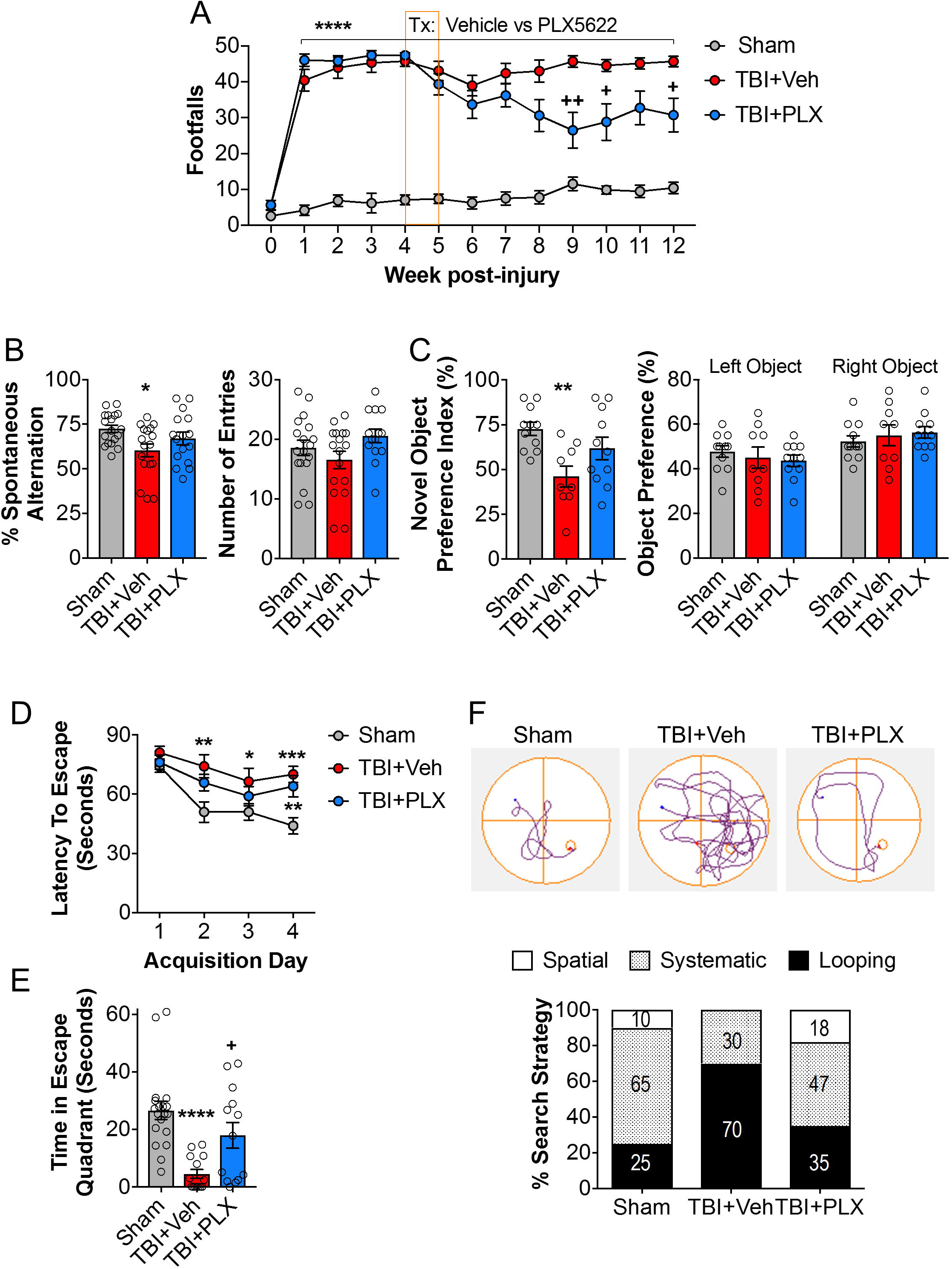
Delayed depletion of microglia starting at 4 weeks post-injury improves long-term motor and cognitive recovery after TBI. Delayed depletion of microglia using the CSF1R inhibitor, PLX5622, significantly reduced long-term TBI-induced fine-motor impairments (number of footfalls in a beam walk task) at 8, 10, and 12 weeks post-injury **(A)**. At 10 weeks post-injury, TBI significantly decreased % spontaneous alterations in the Y-maze task, when compared to sham counterparts. TBI + PLX5622-treated animals had increased % spontaneous alterations, equal to levels in sham **(B)**. At 11 weeks post-injury, TBI significantly decreased % time spent with the novel object in a novel object task, when compared to sham. TBI + PLX5622-treated animals spent increased time with the novel objects, similar to levels of sham **(C)**. In the MWM task at 12 weeks post-injury, TBI + Vehicle- and TBI + PLX5622-treated animals required significantly increased time to locate the hidden submerged escape platform on acquisition days 2-4, when compared to sham **(D)**. There was no significant difference in escape latencies between TBI + Vehicle- and TBI + PLX5622-treated animals. In the probe trial, TBI + Vehicle-treated animals spent significantly less time in the escape quadrant, when compared to sham **(E)**. TBI + PLX5622-treated animals spent significantly more time spent in the escape quadrant, when compared to TBI + Vehicle-treated counterparts. Escape strategy used during the probe trial was assessed, and the percent composition for each strategy (spatial, systematic, and looping) demonstrated that TBI + Vehicle-treated animals used increased looping strategies, and decreased systematic and spatial search strategies when compared to sham and TBI + PLX5622-treated counterparts **(F)**. *p<0.05, ****p<0.0001 vs sham; +p<0.05, ++p<0.01 vs TBI + Vehicle. Data expressed as mean ± s.e.m. (n=9-20/group).

We next investigated cognitive function using a Y-maze task that assesses hippocampal-dependent spatial working memory. When compared to the % spontaneous alternations in sham mice (71.3% ± 2.3%), Vehicle-treated TBI mice had decreased spontaneous alternations in the task, indicative of spatial working memory deficits (60.4 ± 3.6%; TBI effect, F_(2,48)_=3.926; *p*<0.05 vs Sham; **Fig. 1B**). In contrast, PLX5562-treated TBI mice (68.5 ± 3.4) had similar performance as sham mice, indicating improved spatial working memory following PLX5622 treatment. Non-spatial hippocampal-mediated memory was also tested using a novel object recognition task at 11 weeks post-injury. During the familiar phase of the test, there was no difference between sham, Vehicle-treated, or PLX5622-treated TBI mice with regards to time spent with either the right or left objects, indicating no side preference (**Fig. 1C**). Twenty-four hours later, mice were retested with a novel object. When compared to sham mice (78.0 ± 5.3%), Vehicle-treated TBI mice spent significantly less time with the novel object (46.0 ± 6.0%; TBI effect, F_(2,28)_=5.914; *p*<0.01 vs Sham; **Fig. 1C**). PLX5622-treated TBI spent an increased amount of time with the novel object (59.0 ± 7.6) when compared to the Vehicle-treated TBI group; however, differences did not reach statistical significance (**Fig. 1C**).

We performed the Morris Water Maze at 12 weeks post-injury to assess spatial learning and memory. One-way repeated measures ANOVA revealed an effect of time (acquisition day, AD; F_(3,208)_=7.868, *p*<0.001), and PLX5622 treatment (F_(2,208)_=41.63, *p*<0.001), and an interaction between both factors on latency to the escape platform during the test (F_(6, 208)_=2.756, *p*=0.04). When compared to Sham mice, both Vehicle-treated and PLX5622-treated TBI mice had increased latency times during AD2 (*p*<0.01 vs Sham), AD3 (*p*<0.05 vs Sham), and AD4 (*p*<0.001 vs Sham) (**Fig. 1D**). There were no differences in escape latencies between Vehicle-treated and PLX5622-treated TBI mice at any time point. Twenty-four hours later, retention memory was assessed using a probe trial. There was a significant effect of PLX5622 treatment on % time spent in the escape quadrant during the probe trial (F_(2,47)_=11.14, *p*<0.001; **Fig. 1E**). Vehicle-treated TBI mice spent less time in the escape quadrant than sham mice (*p*<0.001 vs Sham). In contrast, PLX5622 treatment reversed TBI-induced retention memory deficits, and PLX5622-treated TBI mice spent more time in the escape quadrant than Vehicle-treated TBI mice (p<0.05 vs TBI+Veh; **Fig. 1E**). There were no differences between groups for swim speeds (**Fig. S5A**), thereby indicating that cognitive improvements were independent of deficits in gross locomotor activity. Finally, to assess search strategies used by mice to find the hidden platform, the swim pattern for each mouse was analyzed and assigned a search strategy (spatial, non-spatial systematic, or repetitive looping; **Fig. S5B**) as previously described (Brody and Holtzman, 2006; Byrnes *et al*., 2012; Zhao *et al*., 2012). Vehicle-treated TBI mice primarily used repetitive looping search strategies (70%) to find the submerged platform when compared to sham mice (25% looping strategy; **Fig. 1F**). In contrast, sham and PLX5622-treated TBI mice predominantly used a systematic search strategy (Sham = 65%; TBI+PLX = 47%) when compared to Vehicle-treated TBI mice (30% systematic search strategy; p<0.001, *x^2^*=38.43; **Fig. 1F**).

### Delayed depletion of microglia using CSF1R inhibitor after TBI reduces histological markers of neurodegeneration and chronic microglial activation

To investigate the effect of temporary depletion of microglia during the chronic stages of TBI on neuropathology, the cortical lesion volume and neuronal loss was assessed at 3 months post-injury. As expected, TBI produced a large cortical lesion in the Vehicle-treated group (7.26 ± 0.76mm^3^; **Fig. 2A**). In contrast, PLX5622 treatment reduced the cortical lesion (5.55 ± 0.52mm^3^; t_(19)_=1.744, *p*<0.05 vs TBI+Veh). Furthermore, TBI induced significant neuronal loss in both the ipsilateral cortex (F_(2,21)_=9.560, *p*<0.001; **Fig. 2C**) and the dentate gyrus (DG) of the hippocampus (F_(2,19)_=6.417, p<0.01; **Fig. 2D**). Post-hoc analysis demonstrated that TBI resulted in chronic neuronal loss in injured cortex (*p*<0.01 vs Sham) and DG (*p*<0.05 vs Sham). Notably, PLX5622-treated TBI mice had reduced neuronal loss in both the cortex (*p*<0.01 vs TBI+Veh; **Fig. 2C**) and DG (*p*<0.05 vs TBI+Veh; **Fig. 2D**).

**Figure 2.**
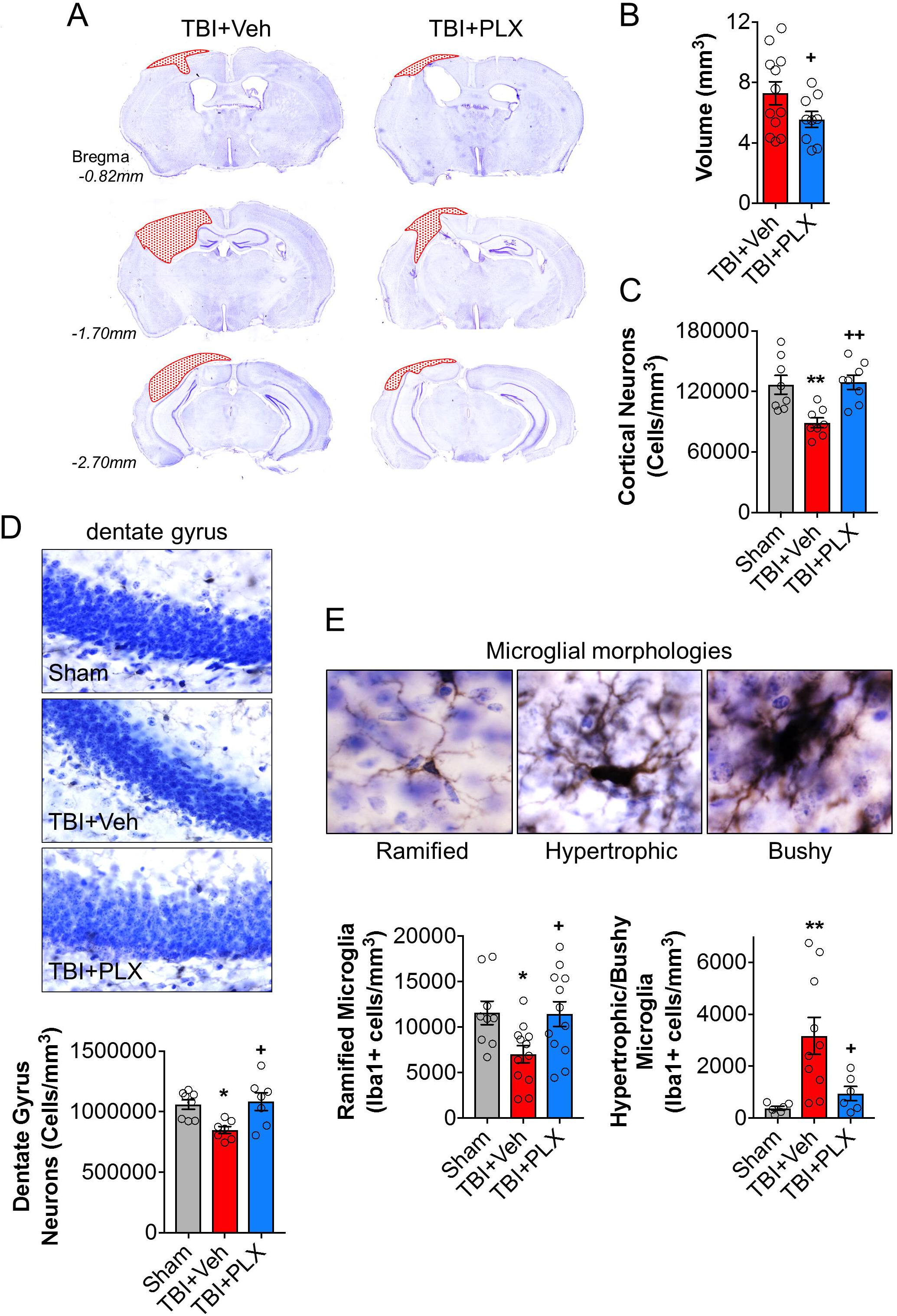
Delayed depletion of microglia starting at 4 weeks post-injury reduces chronic neurodegeneration and alters microglial morphology. Representative images of the cortical lesion in TBI + Vehicle- and TBI + PLX5622-treated animals at 3 months post-injury **(A)**. Stereological analysis demonstrated that TBI + PLX5622-treated animals had a significantly decreased lesion volume, when compared to TBI + Vehicle-treated counterparts **(B)**. Stereological analysis demonstrated that TBI+PLX5622 treatment rescued TBI-induced neuronal loss in the ipsilateral cortex and dentate gyrus (DG) of hippocampus at 3 months post-injury **(C, D)**. TBI significantly reduced cortical neuronal loss, when compared to sham counterparts, whereas PLX56722 treatment prevented this loss such that TBI + PLX5622-treated animals had significantly increased neuronal survival compared to TBI + Vehicle-treated animals **(C)**. Representative images of cresyl violet stained neurons in the DG. TBI significantly reduced DG neuronal loss, when compared to sham. In contrast, TBI + PLX5622-treated animals had significantly increased neuronal survival in DG compared to TBI + Vehicle-treated animals **(D)**. Representative images of Iba1+ microglia displaying ramified, hypertrophic, and bushy morphology phenotypes at 3 months post-injury. Stereological analysis demonstrated that TBI + Vehicle-treated animals had significantly decreased numbers of ramified Iba1+ microglia and increased hypertrophic/bushy Iba1+ microglia in injured cortex, when compared to sham **(E)**. In contrast, TBI + PLX5622-treated animals had significantly increased ramified, and decreased hypertrophic/bushy microglia when compared to TBI + vehicle-treated counterparts. *p<0.05, **p<0.01 vs sham; +p<0.05, ++p<0.01 vs TBI + Vehicle. Data expressed as mean ± s.e.m. (n=8-12/group).

We next performed unbiased stereological assessment of Iba1+ cells to quantify microglial cell number and activation state in the perilesional cortex at 3 months post-injury, as previously described (Byrnes *et al*., 2012; Loane *et al*., 2014). Despite having an equal number of total Iba1+ cells in each group (data not shown), there was an effect of TBI on the number of ramified (F_(2,30)_=4.718, *p*<0.01; **Fig. 2E**) and hypertrophic/bushy microglia (F_(2,19)_=6.884, *p*<0.01; **Fig. 2E**). There were reduced numbers of Iba1+ ramified microglia in the ipsilateral cortex of Vehicle-treated TBI mice compared to sham mice (*p*<0.05; **Fig. 2E**). In contrast, PLX5622 treatment increased numbers of ramified microglia in the injured cortex (*p*<0.05, TBI+PLX vs TBI+Veh; **Fig. 2E**). Similarly, TBI increased the numbers of Iba1+ hypertrophic/bushy microglia in the ipsilateral cortex of Vehicle-treated TBI mice compared to sham mice (p<0.05 vs Sham; **Fig. 2E**). Notably, PLX5622 treatment reduced the numbers of numbers of hypertrophic/bushy Iba1+ microglia in the injured cortex when compared to Vehicle-treated TBI mice (p<0.05, TBI+PLX vs TBI+Veh; **Fig. 2E**).

### Delayed depletion of microglia using CSF1R inhibitor alters cortical transcription patterns related to neuroinflammation, oxidative stress, apoptosis, and neuroplasticity

We then sought to define network-level changes in key components of the cortical transcriptome after TBI by analyzing bulk cortical tissue RNA with Nanostring (Neuropathology panel). Partial least squares discriminant analysis (PLSDA) and hierarchical clustering were used to identify differentially expressed genes between the four treatment groups. PLSDA allows for semi-supervised classification of different treatment groups and determination of signature genes of each experimental variable. In our model, principal component (PC1) captured the variation caused by TBI, while PC2 captured variation caused by PLX5622 treatment (**Fig. 3A**). The PLSDA scores plot shows that gene expression signatures from PCs 1-2 clustered mice from each of the 4 treatment groups as expected: group 1 (Sham+Vehicle), group 2 (TBI+Vehicle), group 3 (Sham+PLX5622), and group 4 (TBI+PLX5622; **Fig. 3A**). The signature genes for each PC are shown in **Figure S6A**, and the loadings indicated the influence of each gene on signatures of TBI (PC1) or PLX (PC2). Genes with more positive loadings on PC1 were upregulated with TBI while negative loadings on PC1 were downregulated by TBI. Overall model performance was very good for all treatment groups (**Fig. S6B**) with classification errors ranging from 84.56% (group 1) to 98.58% (group 4). Group 4 was the least effectively classified due to the large intragroup variance in gene expression, suggesting that TBI+PLX outcomes were more heterogeneous at the level of gene expression. The high classification accuracies for all 4 experimental groups indicated that two PCs captured key signature genes of changes due to TBI and PLX.

**Figure 3.**
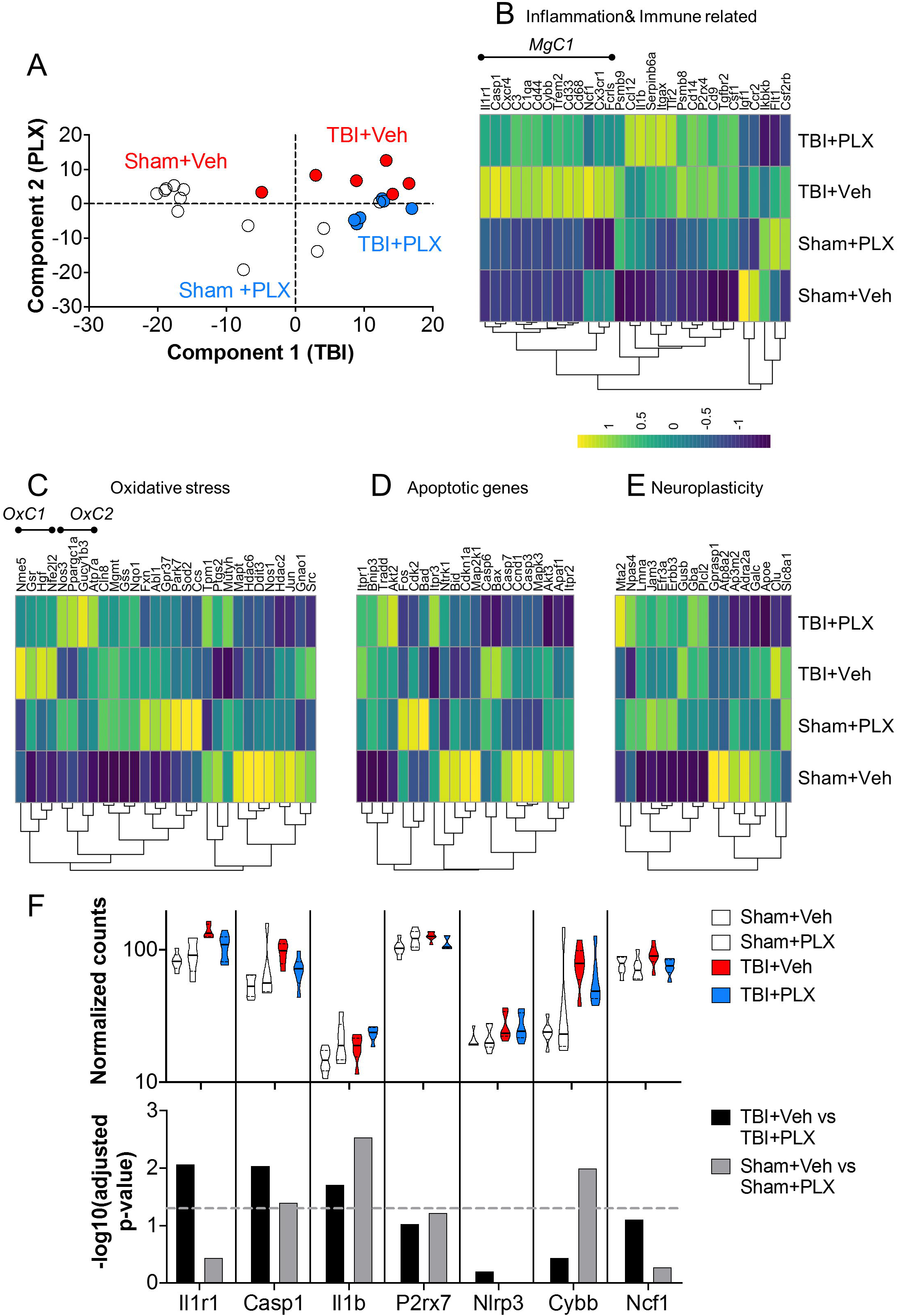
Delayed depletion of microglia during the chronic stages of TBI alters neuroinflammation, oxidative stress, apoptosis, and neuroplasticity pathways in the cortex. Nanostring analysis using a Neuropathology panel was used to assess cortical transcriptional patterns at 2 months post-injury. A partial least squares discriminant analysis (PLSDA) model was generated to classify each of the 4 treatment groups: group 1 (Sham+Vehicle), group 2 (TBI+Vehicle), group 3 (Sham+PLX5622), and group 4 (TBI+PLX5622) **(A)**. Principal components (PC) 1 and 2 identified key genes changed by TBI and PLX, respectively. Hierarchical clustering of differentially expressed genes was performed on different functional annotations. TBI induced a large microglial-related gene cluster (*MgC1*) associated with inflammation and immune function, representing the inflammatory state induced by TBI **(B)**. Analysis of oxidative stress genes identified 2 clusters of genes altered by PLX5622 treatment. The first cluster (*OxC1*) included genes that were increased after TBI, and were significantly reduced by TBI+PLX5622 treatment **(C)**. The other cluster (*OxC2*) was significantly increased in TBI+PLX5622 animals when compared to the TBI + Vehicle animals. Clustering analysis of apoptosis-related genes showed that TBI increased expression of the mitochondrial membrane permeator, Bax, which was decreased by PLX5622 treatment **(D)**. Clustering analysis of neuroplasticity genes showed that the most highly expressed neuroplasticity genes in Sham + Vehicle were downregulated by TBI, including in TBI + PLX5622 treated animals. This indicated that PLX5622 treatment did not return neuroplasticity gene expression to a pre-injured state **(E)**. Analysis of transcript counts (Violin plots and −log10 adjusted P value assessment) for selected genes related to NLRP3 inflammasome (Il1r1, Casp1, Il1b P2rx7, Nlrp3) and NADPH oxidase (Cybb, Ncf1) pathways demonstrated that key genes in both inflammatory pathways were upregulated by TBI, and subsequently reduced by PLX5622 treatment **(F)**. (n=8-9/group).

Differential expression analyses were performed using NanoStringDiff to identify statistically significant differences in gene expression between groups. We focused our analyses on differences in Sham+Vehicle vs Sham+PLX5622 groups, and TBI+Vehicle vs TBI+PLX5622 groups. Analysis of inflammation and immune related genes identified a cluster of microglial-related genes (*MgC1*) that were elevated in TBI+Vehicle compared to Sham+Vehicle and Sham-PLX5622 groups (**Fig. 3B**). Critically, the TBI+PLX5622 group had reduced expression of *MgC1* genes compared to TBI+Vehicle. *MgC1* upregulated genes included inflammatory receptors (e.g., Cd33, Cx3cr1, Cd44 and Cxcr4), the IFNγ-induced immunoproteasome gene Psmb9, and the TGFβ-dependent microglia-specific gene Fcrls. The pro-inflammatory NADPH oxdase-related genes, Cybb and Ncf1, were also up-regulated in TBI+Vehicle group, and attenuated in the TBI+PLX5622 group. We next examined 2 clusters of genes involved in oxidative stress responses that were changed by PLX5622 treatment (**Fig. 3C**). The first cluster (*OxC1*) included genes upregulated by TBI, with highest expression in TBI+Vehicle group that was decreased by PLX5622 treatment (e.g., Nme5, Gsr, Hgf, Hfe2l2). Another cluster (*OxC2*) of genes (Ppargc1a, Nos3, Gucy1b3, Atp7a) demonstrated an inverse trend: they had increased expression in the TBI+PLX5622 group when compared to the TBI+Vehicle group. Apoptosis-related genes were also evaluated (**Fig. 3D**). Broadly, each experimental group had a distinct pattern of apoptotic gene expression. PLX5622 acted partially by reducing expression of key caspases (Casp6, Casp7) in injured cortex. Treatment with PLX5622 also decreased the upstream pro-apoptotic genes Bax, a mitochondrial membrane permeator. Finally, neuroplasticity genes were also assessed (**Fig. 3E**). The most highly expressed genes in Sham + Veh were most downregulated in TBI + PLX, indicating that PLX treatment does not return neuroplasticity gene expression to a pre-injured state. This result mirrored expression patterns in apoptotic genes.

Focused analysis of transcript counts for selected genes related to NLRP3 inflammasome (Il1r1, Casp1, P2rx7, Nlrp3) and NADPH oxidase (Cybb, Ncf1) pathways demonstrated that key genes in both inflammatory pathways are reduced by PLX5622 treatment during chronic TBI (**Fig. 3F**). The up-regulation of Il1b mRNA with PLX5622 treatment may reflect a compensatory response. Therefore, network-level gene changes identified by Nanostring analysis suggested that PLX5622 treatment acted through multiple beneficial mechanisms to improve chronic TBI outcomes, including reconditioning cells to be less pro-inflammatory, reducing oxidative stress sensing and responses and altering expression of key NADPH oxidase and NLRP3 inflammasome components.

Finally, we confirmed a subset of microglial-related inflammatory pathways in injured cortex using real-time PCR and immunofluorescence imaging. Short-term depletion and subsequent repopulation of microglia during the chronic stages of TBI by PLX5622 treatment decreased TBI-induced upregulation of NLRP3 inflammasome related genes (Nlrp3, Casp1, and Il1b; **Fig. 4A**), and NADPH oxidase genes (Cybb [NOX2], Cyba [p22^phox^], Ncf1, Ncf4; **Fig. 4B**) in bulk cortical RNA. Immunofluorescence imaging confirmed NADPH oxidase (NOX2+) expression in reactive microglia (Iba1+/CD68+) in the injured cortex at 3 months post-injury (**Fig. 4C**). Notably, the number of NOX2+/CD68+ reactive microglia was reduced in the injured cortex of PLX5622-treated TBI mice (t_(11)_ = 2.189; *p*<0.05 vs TBI+Veh; **Fig. 4C**).

**Figure 4.**
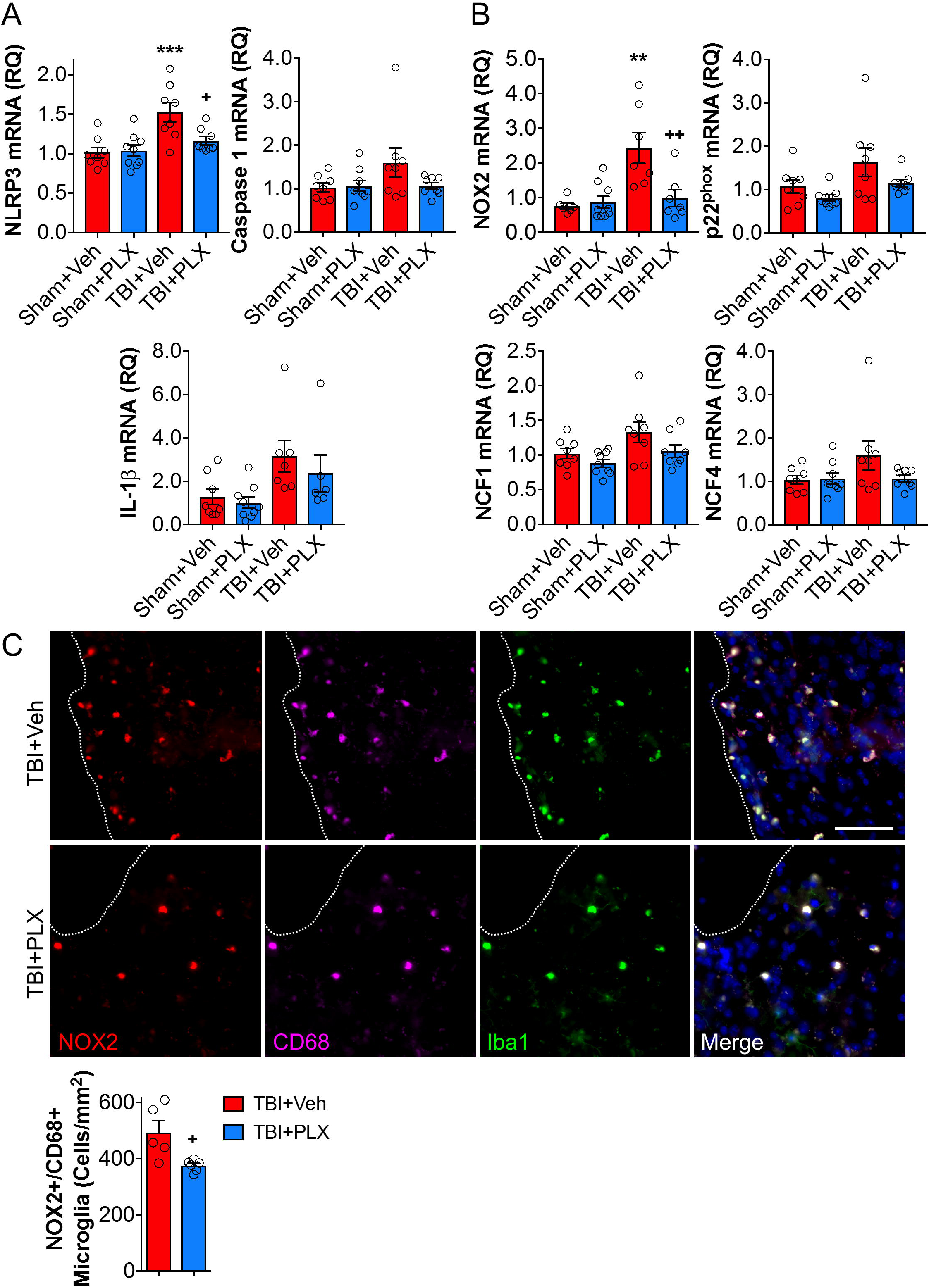
Delayed depletion of microglia decreases chronic NLRP3 inflammasome and NOX2 expression in the injured cortex. Analysis of ipsilateral cortical tissue at 2 months post-injury demonstrated that TBI increased Nlrp3, Casp1, and Il1b mRNA. The TBI-induced increase in Nlrp3 mRNA was significantly reduced by PLX5622 treatment **(A)**. TBI increased NADPH oxidase subunits, Cybb (NOX2), Cyba (p22^phox^), Ncf1, and Ncf44 mRNA in injured cortex. The TBI-induced increase in Cybb mRNA was significantly reduced by PLX5622 treatment **(B)**. Immunofluorescence analysis of reactive microglia (NOX2+, red; CD68+, cyan; Iba1+, green) in the injured cortex at 3 months post-injury demonstrates that delayed depletion of microglia significantly decreased the number of NOX2+ reactive microglia when compared to levels in TBI + Vehicle-treated animals **(C)**. Scale bar = 100μm. *p<0.05, **p<0.01, ***p<0.001 vs sham animals; +p<0.05, ++p<0.01 vs TBI + Vehicle-treated animals. Data expressed as mean ± s.e.m. (n=6-9/group)

### Delayed depletion of microglia using CSF1R inhibitor reduced the inflammatory status of microglia at 2 months post-injury

To better understand the cellular correlates related to chronic transcriptional changes in the cortex, we evaluated inflammatory responses in microglia using flow cytometry. TBI increased absolute numbers of live CD45^int^/CD11b^+^/Ly6C^-^ microglia at 2 months post-injury when compared to sham levels (**Fig. 5A**); PLX6522 treatment had no effect on microglial cell numbers at this time point. The total number of infiltrating CD45^hi^ leukocytes were also increased by TBI when compared to sham levels (**Fig. 5A**), but relative numbers were low (291 ± 44 CD45^hi^ cells) compared to numbers of resident microglia (48474 ± 3108 CD45^int^ cells). Again, PLX6522 treatment had no effect on CD45^hi^ cell numbers. Microglial morphological features were examined including analysis of microglial size by forward scatter (FSC) and microglial cellular granularity by side scatter (SSC). Consistent with histological findings for hypertrophic/bushy Iba1+ microglia (**Fig. 2E**), TBI increased microglial cell size (FSC+; F_(2,15)_=10.02, p<0.05 vs Sham) and granularity (SSC+; F_(2,19)_=21.34, p<0.05 vs Sham) when compared to levels in sham mice (**Fig. 5B**). PLX5622 treatment reduced both size and granularity parameters in microglia (FSC+: p<0.01 vs TBI+Veh; SSC+: p<0.001 vs TBI+Veh). Assembly of NLRP3 inflammasome leads to caspase-1-dependent release of pro-inflammatory IL-1β (Heneka *et al*., 2018), so we assessed caspase-1 activity and IL-1β expression in microglia. Consistent with mRNA analysis (**Fig. 4A**), TBI increased caspase-1 activity (F_(2,10)_=5.608, p<0.05 vs Sham; **Fig. 5C**) and IL-1β production (F_(2,11)_=14.01, p<0.001 vs Sham; **Fig. 5D**) in microglia when compared to levels in sham mice. Notably, PLX5622 treatment reduced microglial caspase-1 activity and IL-1β production (p<0.01 vs TBI+Veh). Therefore, delayed depletion of microglia using PLX5622 attenuates key markers of NLRP3 inflammasome activation in microglia during chronic stages of TBI.

**Figure 5.**
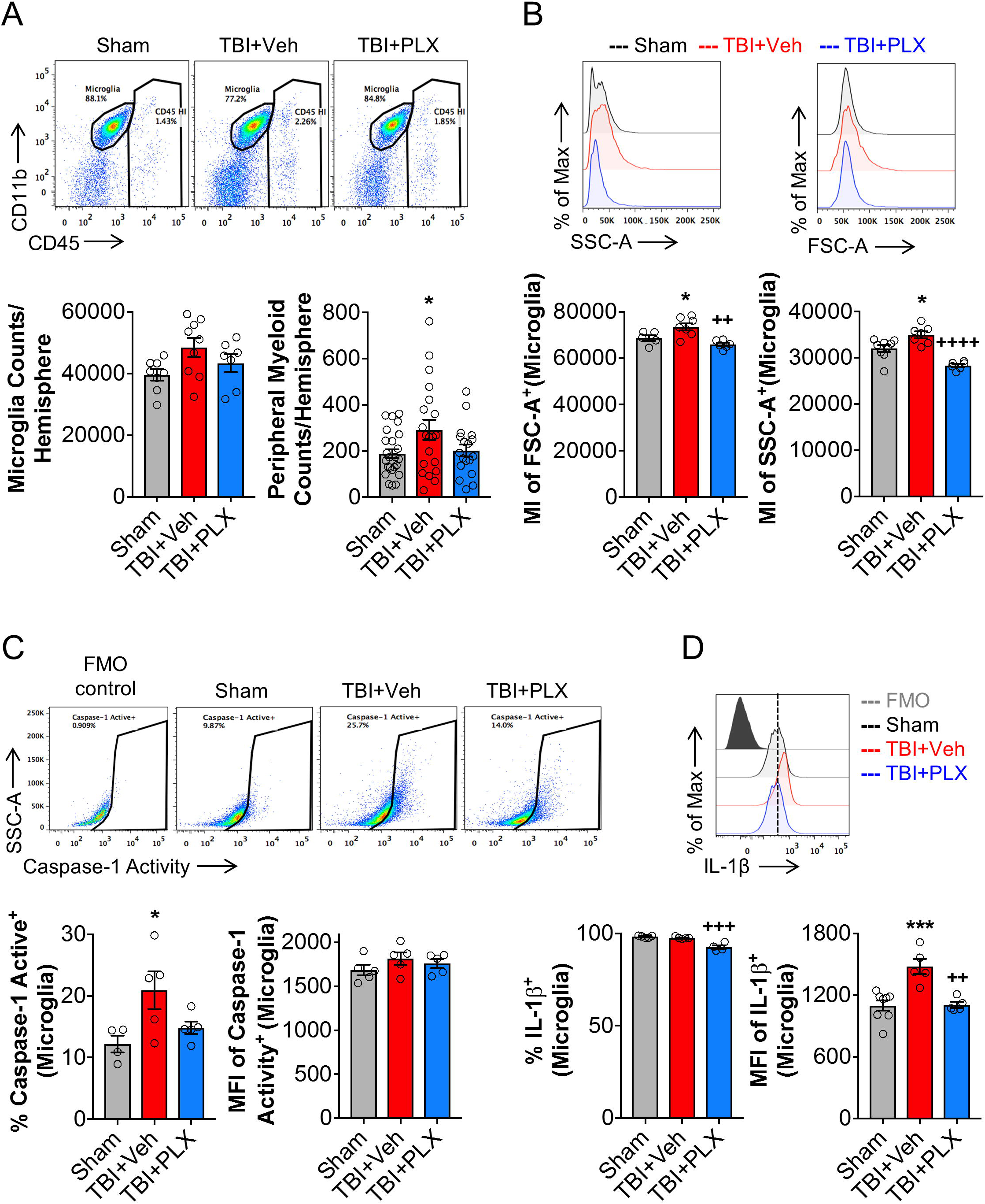
Delayed depletion of microglia reduced chronic expression of NLRP3 inflammasome components in microglia. Flow cytometry analysis and representative dot plots show populations of resident CD45^int^ microglia and infiltrating peripherally derived CD45^hi^ myeloid cells in the ipsilateral cortex at 2 months post-injury **(A)**. There was no difference in absolute microglial numbers between sham, TBI + Vehicle-treated, and TBI + PLX5622-treated groups. TBI resulted in a significant increase in CD45^hi^ myeloid cells in brain in TBI + Vehicle-treated animals, but numbers were small in comparison to numbers of resident microglia in brain at this timepoint. Assessment of microglial cell size as measured by forward scatter (FSC), and microglial granularity as measured by side scatter (SSC), demonstrated that TBI increased microglial cell size and granularity when compared to levels in sham microglia **(B)**. PLX5622 treatment significantly reduced both markers in microglia. Representative dot plots show resident CD45^int^ microglia expressing active caspase 1. TBI significantly increased active caspase 1 in microglia, when compared to levels in sham **(C)**. Levels of active caspase 1 were decreased by PLX5622 treatment, but failed to reach statistical significance. TBI also significantly increased IL-1β in microglia, when compared to levels in sham. Levels of microglial IL-1β was significantly decreased in TBI + PLX5622-treated animals **(C, D).** *p<0.05, ***p<0.001, vs Sham; ++p<0.01, +++p<0.001, ++++p=0.000 vs TBI + Vehicle-treated animals. Data expressed as mean ± s.e.m. (n=5-9/group)

### Delayed depletion of microglia using CSF1R inhibitor does not affect post-traumatic astrocytic responses

In response to TBI, astrocytes tune their reactivity to varying degrees of axonal injury, vascular disruption, ischemia, and inflammation (Burda *et al*., 2016). Astrocytes and microglia dynamically communicate via cytokines and cell-surface markers during aging, neurodegeneration and CNS injury (Villacampa *et al*., 2015; Liddelow *et al*., 2017). Therefore, to determine the effect of delayed PLX5622 treatment after TBI on astrocytes in injured cortex we evaluated GFAP expression, a canonical marker of astrocytes. TBI increased GFAP mRNA expression in the cortex (F_(2,19)_=4.644, *p*<0.05) when compared to levels in sham mice (p<0.05 vs Sham; **Fig. S7A**). GFAP mRNA was decreased by PLX5622 treatment, but differences compared to the Vehcile-treated TBI group did not reach statistical significance. We also performed *in situ* analysis to evaluate astrocyte populations in the perilesional region at 3 months post-injury. TBI increased astrocyte cellular area (F_(1,18)_=29.99, *p*<0.05; **Fig. S7Bi, C**) and GFAP expression (F_(1,18)_=21.36, *p*<0.01; **Fig. S7Bii, C**). PLX5622 treatment did not change post-traumatic GFAP cellular distribution or expression at 3 months post-injury (**Fig. S7Bi, C**).

## Discussion

Recent clinical and experimental studies have revised the outdated concept that TBI is an acute neurological disorder (Wilson *et al*., 2017). Instead, persistent and diffuse posttraumatic neuroinflammation, with progressive neurodegeneration, can continue for months to years in TBI patients (Gentleman *et al*., 2004; Ramlackhansingh *et al*., 2011; Johnson *et al*., 2013; Coughlin *et al*., 2015; Cherry *et al*., 2016; Coughlin *et al*., 2017). Microglia undergo changes in their activation profile during the chronic inflammatory response that appears to contribute to cognitive decline in many neurodegenerative disorders (Perry *et al*., 2010; Heneka *et al*., 2015). Epidemiological studies show that a prior history of TBI is associated with an increased incidence of both Alzheimer’s disease (AD) and non-AD forms of dementia (Mortimer *et al*., 1985; Dams-O’Connor *et al*., 2013; Gardner *et al*., 2014; Nordstrom *et al*., 2014; Wilson *et al*., 2017), with the latter appearing to be most prevalent. Experimental evidence suggests that such neurodegeneration is associated with maladaptive transformation of microglia from a neurorestorative and neuroprotective phenotype to a persistent, dysfunctional neurotoxic phenotype (Faden and Loane, 2015; Simon *et al*., 2017). Thus, chronic release of neurotoxic molecules, including reactive oxygen species, and nitric oxide, as well as pro-inflammatory cytokines, from reactive microglia may cause chronic progressive neuronal loss and/or white matter degeneration over months to years (Faden and Loane, 2015; Simon *et al*., 2017).

The present study demonstrates that short-term depletion and subsequent repopulation of microglia during the chronic stages of experimental TBI reduces chronic neuroinflammation, improves neurological recovery (sensori-motor and cognitive), and decreases neurodegeneration. These findings provide further evidence that the therapeutic window for targeting chronic neuroinflammation and improving functional recovery following TBI may be far longer than traditionally believed (i.e. hours post trauma) (Faden *et al*., 2016). One-week treatment with the CSF1R inhibitor, PLX5622, beginning 4 weeks after TBI removed neurotoxic microglia from the chronically injured brain, and repopulated microglia had an altered phenotype that was markedly less pro-inflammatory. Delayed short-term microglial depletion significantly improved neurobehavioral recovery through 3 months post-injury, as demonstrated by a variety of complementary motor and cognitive tests. Improved functional outcomes were associated with decreased cortical lesion volume and neuronal cell loss in both the cortex and the hippocampus. Furthermore, stereological assessments demonstrated that repopulated microglia during the chronic stages of TBI displayed a more ramified morphology, similar to that of uninjured sham controls, and in contrast to untreated injured animals that displayed the typical chronic posttraumatic hypertrophic morphology (Loane *et al*., 2014). PLX5622 treated animals also had reduced expression of neuroinflammatory and oxidative stress markers in injured cortex, significantly fewer NOX2-positive microglia, and reduced microglial caspase-1 activity and IL-1β production (both components of the NLRP3 inflammasome), indicative of a less pro-inflammatory and reactive microglial phenotype. Thus, short-term depletion of microglia relatively late after TBI largely eliminates the destructive neurotoxic microglial phenotype that appears to contribute importantly to chronic posttraumatic neurodegeneration and associated neurological dysfunction.

CSF1R is vital for the development and survival of microglia (Ginhoux *et al*., 2010; Elmore *et al*., 2014). Novel CSF1R inhibitors have recently been used in experimental models to investigate microglial dynamics during physiological and pathophysiological conditions (Han *et al*., 2019). The latest generation inhibitor, PLX5622, is highly selective for the CSF1R, has desirable pharmacokinetics across multiple species, including oral bioavailability and a >20% brain penetrance, and can produce rapid and sustained elimination of microglia (>95% elimination) (Spangenberg *et al*., 2019). Although microglia play an important role in CNS homeostasis (Salter and Stevens, 2017), depletion of microglia using first generation CSF1R inhibitors (PLX3397) did not result in cognitive or motor impairments in non-injured mice (Elmore *et al*., 2014), and animals with repopulated microglia performed similar to control non-depleted animals in complex cognitive, motor and affective behavioral tasks (Elmore *et al*., 2015). Consistently, we found that depletion and subsequent repopulation of microglia, using PLX5562, did not cause neurobehavioral changes in sham mice. Remarkably, short term depletion of microglia late after TBI markedly reduced TBI-induced impairments in sensori-motor function up to 3 months post-injury, exemplified by decreased numbers of footfalls on the beam walk task and reduced disability on the rotorod test. In addition, when compared to non-treated TBI mice, PLX5622-treated TBI mice had improved hippocampal spatial and working memory, as shown by improved performance in the Morris water maze, Y-maze, and novel object recognition tests. The observed improvements in cognitive function recovery with the delayed PLX5622 treatment paradigm mimics long-term recovery trajectories promoted by first generation CSF1R inhibitors in a mouse hippocampal lesion model (Rice *et al*., 2015).

Previous work from our laboratory demonstrated that experimental TBI in mice induces chronic microglial activation up to one year post-injury and contributes to progressive lesion expansion, hippocampal neurodegeneration, and white matter damage (Loane *et al*., 2014). NADPH oxidase (NOX2) has been implicated as a common and essential mechanism of microglia-mediated neurotoxicity (Babior, 1999; Qin *et al*., 2004; Block *et al*., 2007). We reported that inhibiting NOX2 activity after TBI in mice suppresses microglial neurotoxicity, resulting in reduced chronic tissue loss and improved long-term neurological recovery (Kumar and Loane, 2012; Loane *et al*., 2013; Kumar *et al*., 2016b; Barrett *et al*., 2017). Others have shown that NOX2 is upstream of the NLRP3 inflammasome, and that inhibition of NOX2 reduces post-traumatic neuroinflammation by regulating NLRP3 inflammasome (Ma *et al*., 2017). Previously, we found that pharmacological inhibition of chronic microglial activation beginning 1-month post-injury using either mGluR5 agonists or delayed aerobic exercise, reduced neurodegeneration and improves long-term motor and cognitive recovery (Byrnes *et al*., 2012; Piao *et al*., 2013). Thus, these, and other pre-clinical studies (Ferguson *et al*., 2017), provide evidence for an expanded therapeutic window for neuroprotection by targeting chronic post-traumatic neuroinflammatory responses.

Given that activation of NLRP3 inflammasome has been implicated in chronic neurodegeneration (Heneka *et al*., 2018) and TBI pathophysiology in humans (Adamczak *et al*., 2012; Kerr *et al*., 2018) and rodents (Liu *et al*., 2013; Xu *et al*., 2018; Kuwar *et al*., 2019), and that NOX2 may be an important upstream regulator of NLRP3 inflammasome activation (Dostert *et al*., 2008), we also examined molecular and cellular markers of this pathway. Key genes involved in NLRP3 inflammasome activation and function were chronically upregulated after TBI and reduced by PLX treatment - including Il1r1 and Casp1. Importantly, the inflammatory profile of repopulated microglia at 2 months post-injury showed decreased microglial IL-1β and caspase-1 activation. Ma et al recently reported robust activation of NLRP3 inflammasome in the injured cortex following experimental TBI in mice, and that NOX2 inhibition markedly attenuated its activity (Ma *et al*., 2017). Whereas Ma et al., reported changes during the subacute phase post-TBI (14 days post-injury (Ma *et al*., 2017)), we demonstrated chronic NLRP3 activation (active caspase-1 and IL-1β expression) in microglia at 2 months post-injury, and that PLX5622 treatment decreased NLRP3 inflammasome component mRNA expression in the hippocampus. Thus, PLX5622 treatment confers neuroprotection after TBI, in part, by removing chronic NOX2-associated NLRP3 inflammasome activation in microglia.

To assess the effect of depletion and subsequent microglial repopulation on other glial cells in the CNS, we examined gene and protein expression of GFAP, a marker of astrocyte activation. Astrocyte reactivity depends on varying degrees of axonal injury, vascular disruption, ischemia and inflammation (Burda *et al*., 2016). Repopulated microglia did not significantly affect the TBI-induced increase in GFAP expression or reactivity (morphology) in perilesional cortex at 3 months post-injury. Although previous data regarding the effects of microglial depletion on acute GFAP changes after TBI are conflicting (Witcher *et al*., 2018), we did not observe significant changes in GFAP expression or morphology at chronic timepoints after TBI. Reported differences of PLX5622 treatment on posttraumatic astrocyte reactivity may reflect differences in the injury model (midline fluid percussion vs controlled cortical injury), time-points examined (3 days vs 3 months post-injury), or the treatment paradigm employed (pre-treatment vs late posttrauma treatment). In addition, Witcher et al. reported that depletion of microglia prior to TBI did not alter injury-induced neuronal injury in the cortex at 7 days post-injury (Witcher *et al*., 2018). In contrast, our study demonstrates that delayed depletion of microglia during the chronic stages of TBI significantly reduce neuronal cell loss in cortex and hippocampus. Again, such differences may reflect treatment timing or model.

Taken together, our preclinical studies strongly support the concept that sustained pro-inflammatory and neurotoxic microglial activation contributes to posttraumatic neurodegeneration and related neurological dysfunction after TBI. The neuroprotective effects of short-term depletion of microglia during the chronic stages of TBI likely reflect multi-factorial mechanisms-including less NOX2 mediated neurotoxic inflammation associated with altered oxidative stress responses, reduced apoptotic potential, and altered expression of the NLRP3 inflammasome. These studies support new concepts that the therapeutic window for TBI may be far longer than traditionally believed if chronic and evolving neuroinflammation can be inhibited or regulated in a precise manner (Faden and Loane, 2015). These observations may have important therapeutic implications for human TBI, and suggest that CSF1R inhibition may be a clinically feasible approach to limit posttraumatic neurodegeneration and neurological dysfunction following head injury.

## Abreviations

TBI: traumatic brain injury
CCI: controlled cortical impact
CSF1R: colony stimulating factor 1 receptor
PLX5622: Plexxikon 5622
NOX2: NADPH oxidase
NOR: novel object recognition
MWM: morris water maze
DG: dentate gryus

## Acknowledgements

The authors thank Victoria Meadows and Wesley Shoap for help with neurobehavioral assessment and histology. The authors thank Plexxikon Inc. for the use of PLX5622. This work was supported by National Institutes of Health grants R01NS082308 (D.J.L), R01NS037313 (A.I.F), R01NS096002 (B.A.S), and R01NS110756 (D.J.L./A.I.F/B.A.S), R21EY029451 (J.B.L), a U.S. Veterans Affairs grant 1I01 RX001993 (B.A.S), and Science Foundation Ireland grant 17/FRL/4860 (D.J.L).

## Author contributions

Rebecca J. Henry contributed to study design, performed *in vivo* studies, collected data, performed data analysis, and manuscript preparation; Rodney M. Ritzel contributed with experimental design, collected and performed data analysis of flow cytometry; James P. Barrett contributed to data collection and PCR analysis; Sarah J. Doran contributed to data collection and IHC analysis; Yun Jiao performed nanostring data analysis; Jennie B. Leach contributed to nanostring data analysis; Gregory L. Szeto contributed to nanostring data analysis and manuscript preparation; Bogdan A. Stoica contributed with experimental design; Alan I. Faden contributed with manuscript preparation; David J. Loane contributed to study conception and design, and manuscript preparation. All authors read and approved the manuscript prior to submission.

---

**Supplementary Figure 1. Oral administration of the CSF1R inhibitor, PLX5622, resulted in depletion of microglia in mouse brain.** Representative immunofluorescence images of P2Y12+ microglia demonstrated that oral administration of PLX5622 (1200 ppm) in chow for 7 days leads to an almost complete depletion of microglia (P2Y12+, red) in the mouse brain **(A)**. Scale bar = 200μm. Flow cytometry analysis of microglia (CD11b+/CD45^int^) demonstrated that 7-day PLX5622 treatment resulted in approximately 95% depletion of microglia counts in the mouse brain. Returning animals to normal chow feed resulted in repopulation of microglia in mouse brain within 7 days of removing PLX5622 chow **(B)**. Data expressed as mean ± s.e.m. (n=3-4/group).

**Supplementary Figure 2. Experimental timeline.** Adult male C57Bl/6 mice underwent CCI/Sham surgery. At 4 weeks following CCI/Sham surgery mice were placed on PLX5622 (1200 ppm) or normal chow (Vehicle) for 1 week, following which mice were returned to normal chow for the remainder of the study. One cohort of mice were sacrificed at 8 weeks post-injury and samples were collected for flow cytometry and nanostring analysis. A separate cohort of mice underwent a battery of neurobehavioral tasks (beam walk, Y maze, Novel object recognition, Morris water maze) through 12 weeks post-injury, and samples were collected for histological analysis (Stereology).

**Supplementary Figure 3. Delayed depletion of microglia in sham animals does not result in neurological impairments.** Sham + PLX5622-treated animals did not have fine motor coordination deficits and performed at a similar level as Sham + Vehicle-treated animals in the beam walk task **(A)**. Similarly, Sham + PLX5622-treated animals did not have deficits in cognitive function exemplified by no deficits in % spontaneous alterations in the Y-maze task **(B)**, preference for the novel object in the NOR **(C)**, and latency to the escape platform **(D)**, and time in the escape quadrant in the MWM task **(E)**, when compared to Sham + Vehicle-treated animals. Data expressed as mean ± s.e.m. (n=6-10/group).

**Supplementary Figure 4. Delayed depletion of microglia starting at 4 weeks post-injury improves motor performance in an accelerating rotarod after TBI.** At 12 weeks post-injury, TBI + Vehicle-treated animals spent less time on the accelerating rod when compared to sham counterparts. TBI + PLX5622-treated animals spent increased time on the rotarod, equal to levels in sham; however, differences were not statistically significant. Data expressed as mean ± s.e.m. (n=7-9/group).

**Supplementary Figure 5. Swim speed and search strategy criteria for the Morris water maze.** TBI + Vehicle-treated and TBI + PLX5622-treated animals had similar swim speeds to that of sham operated animals during the acquisition days of the Morris water maze (MWM) **(A)**. Representative images of search strategies (spatial, non-spatial systemic, and repetitive looping) used by animals during the probe trial **(B)**. (n=15-20/group).

**Supplementary Figure 6. PLSDA model gene loadings and performance.** Plot of the top 20 upregulated (blue) and downregulated (red) genes for PC1 and 2 **(A)**. The loadings indicated the influence of each gene on signatures of TBI (PC1) or PLX (PC2). Genes with more positive loadings on PC1 were upregulated with TBI, while negative loadings on PC1 were downregulated by TBI. Genes with negative loadings on PC2 were upregulated in PLX, while positive loadings were downregulated. Classification performance per group was determined by bootstrap resampling. Receiver operating characteristic (ROC) curves showed overall model performance for each treatment group with average performance reported **(B)**.

**Supplementary Figure 7. Delayed depletion of microglia does not alter astrocyte reactivity in the cortex after TBI.** Analysis of ipsilateral cortical tissue at 2 months post-injury demonstrated that TBI increased Gfap mRNA. The TBI-induced increase in Gfap mRNA was not altered by PLX5622 treatment **(A)**. *In situ* analysis of astrocyte populations in the perilesional region at 3 months post-injury. TBI increased astrocyte cellular area and GFAP expression when compared to levels in sham counterpart. PLX5622 treatment did not change post-traumatic GFAP cellular distribution or expression **(B)**. Representative images of GFAP (green) in the perilesional cortex at 3 months post injury **(C)**. Scale bar = 100μm. *p<0.05, **p<0.01 vs sham. Data expressed as mean ± s.e.m. (n=5-8/group).

