## Supplementary figures and images for "Microglial depletion with CSF1R inhibitor during chronic phase of experimental traumatic brain injury reduces neurodegeneration and neurological deficits"

### Supplemental Figure 1

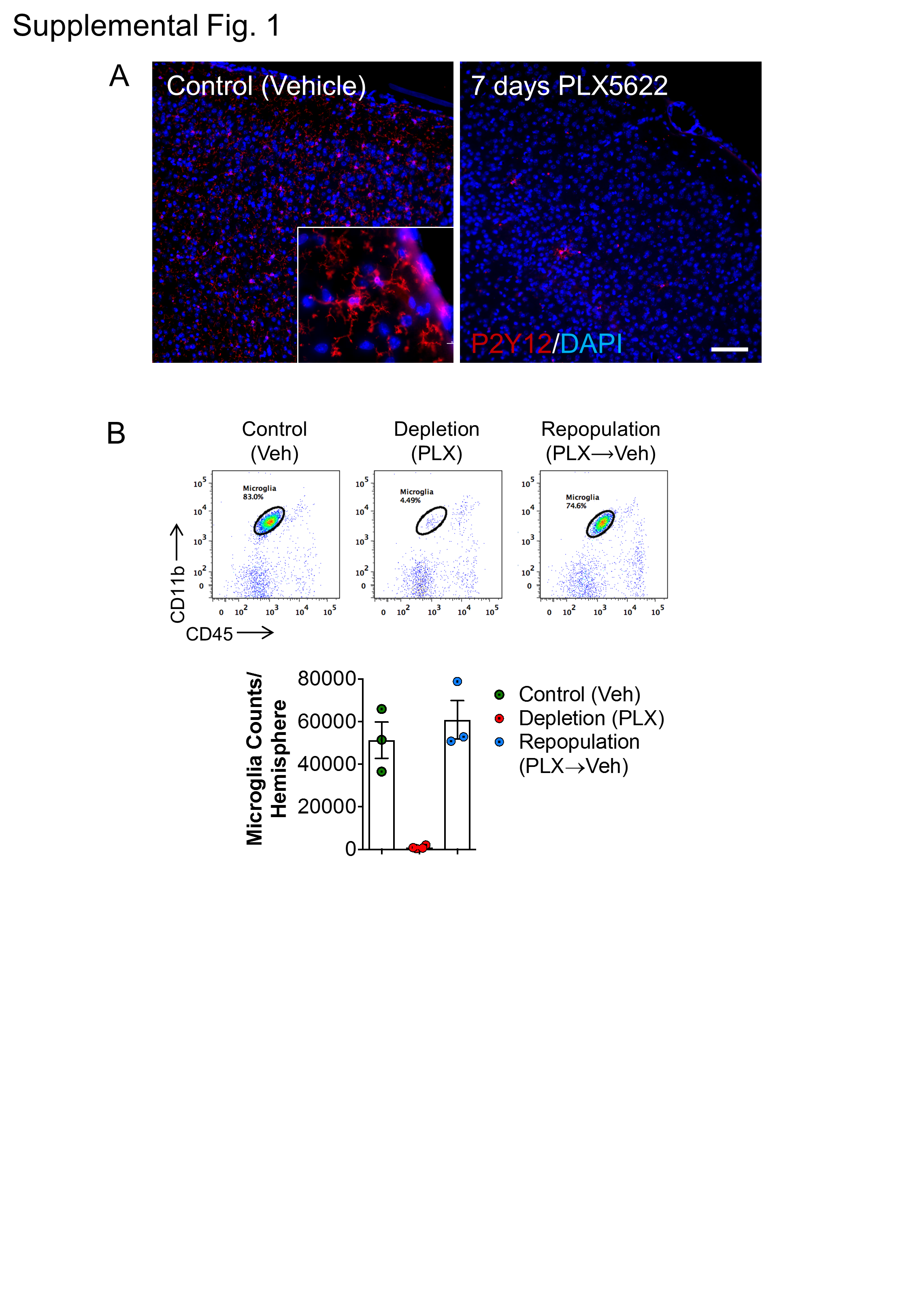

### Supplemental Figure 2

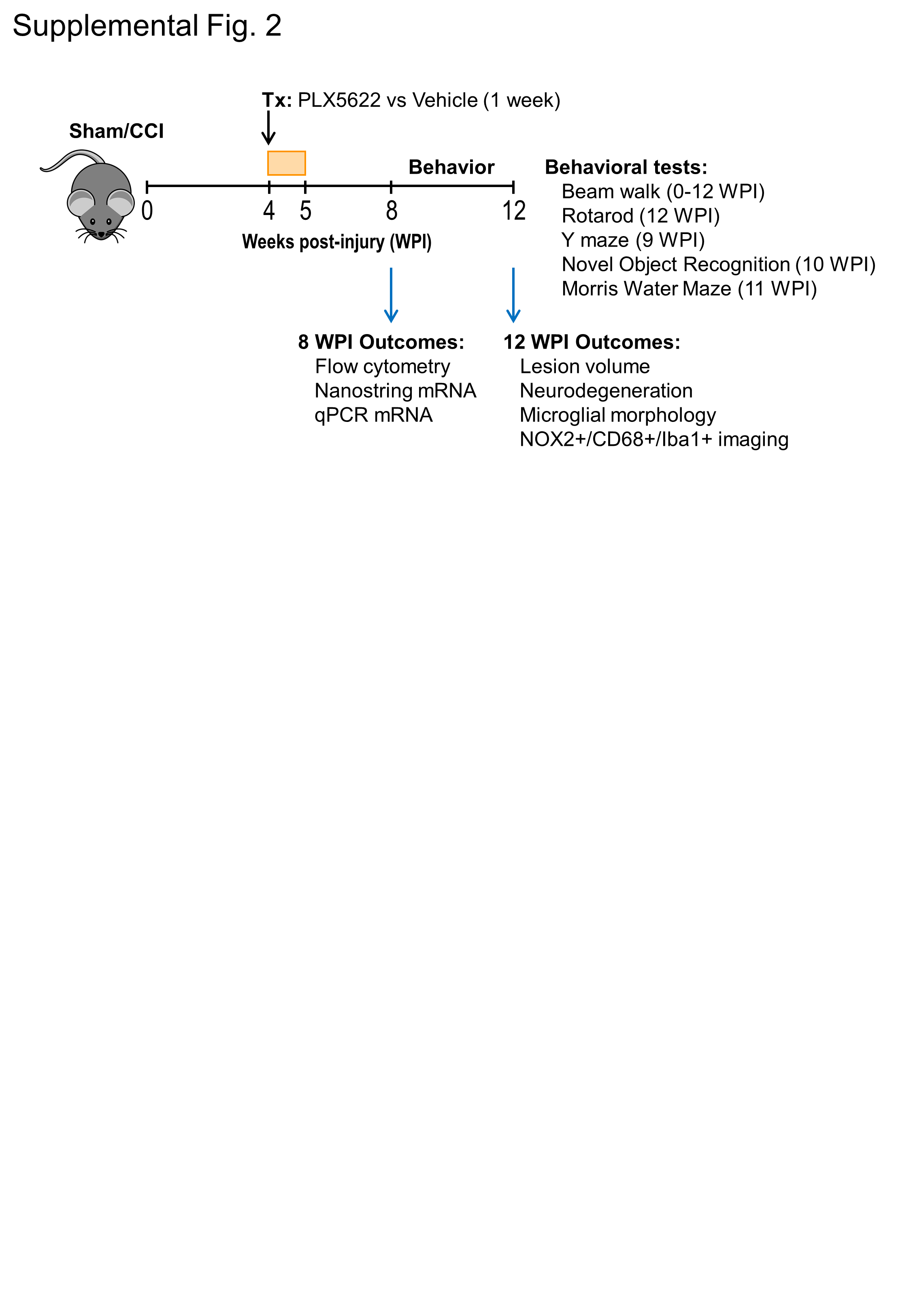

### Supplemental Figure 3

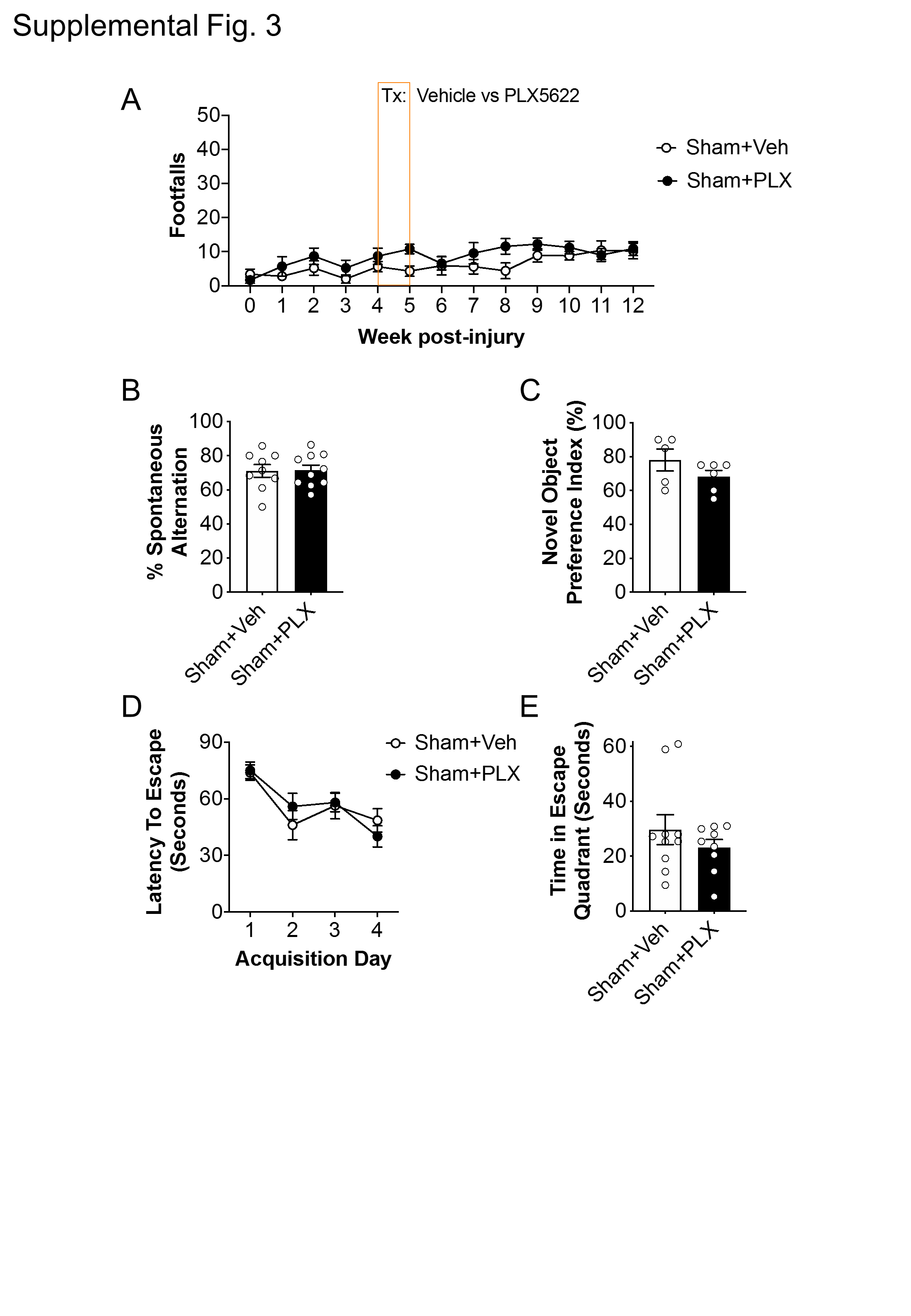

### Supplemental Figure 4

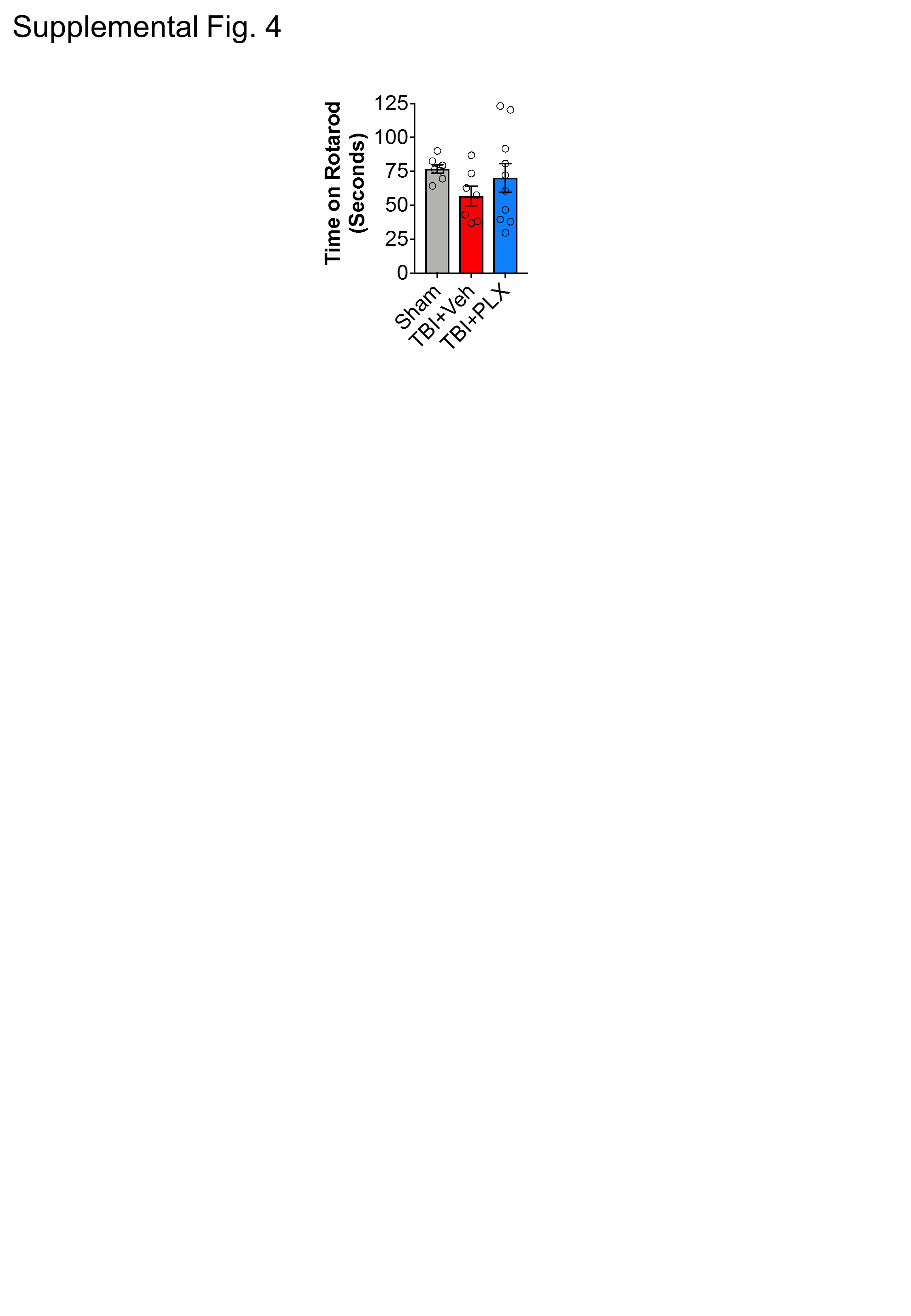

### Supplemental Figure 5

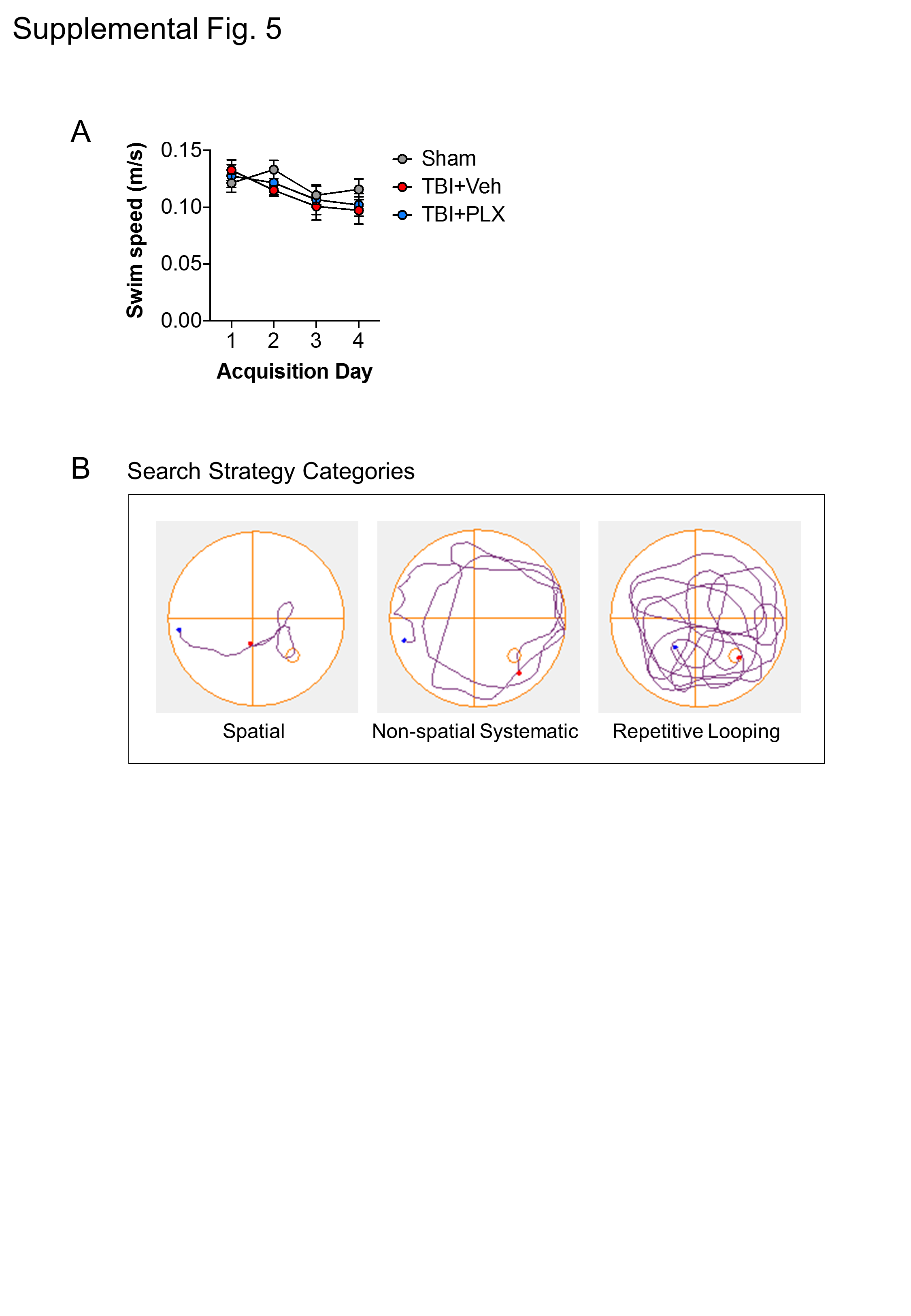

### Supplemental Figure 6

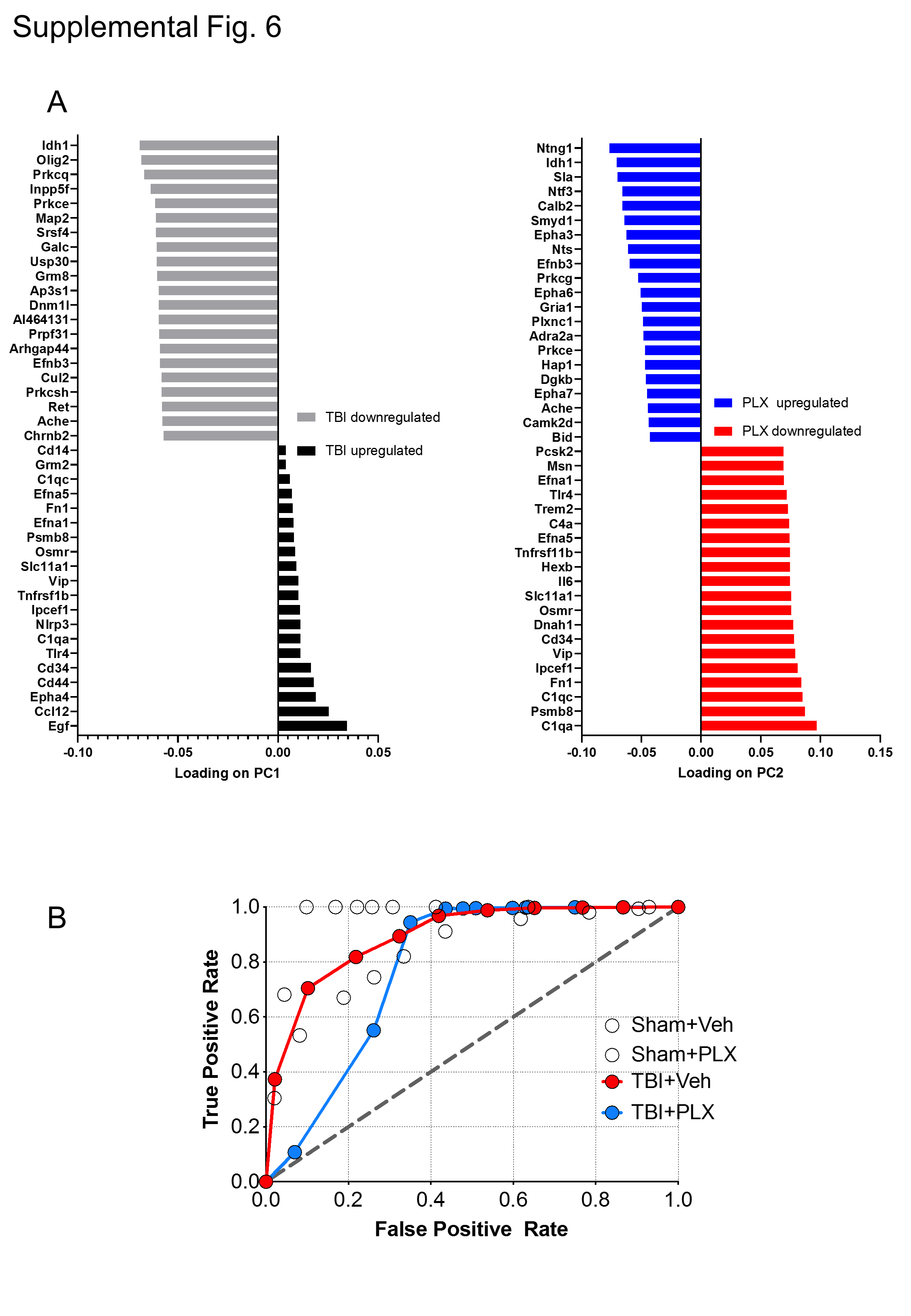

### Supplemental Figure 7

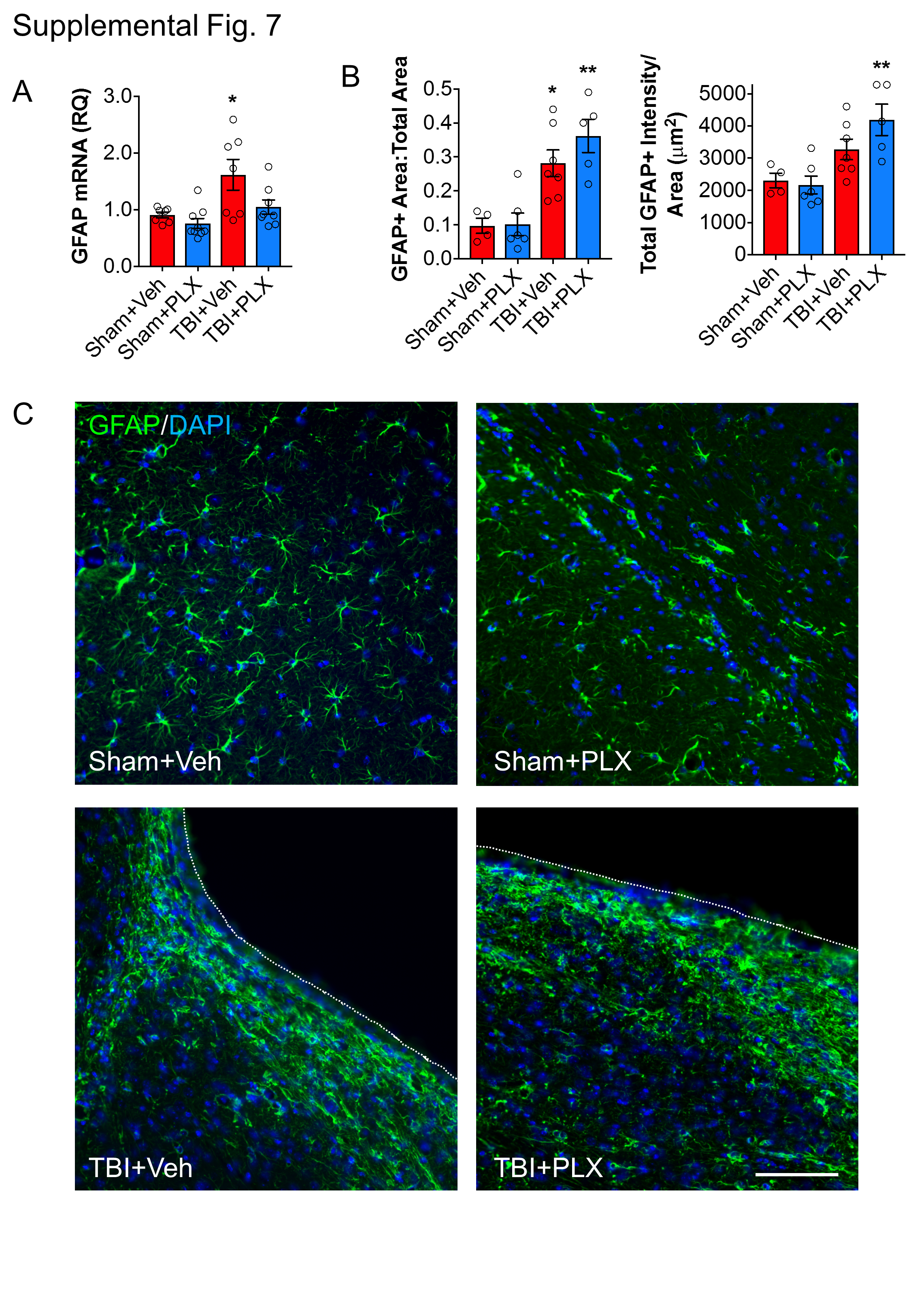
